# *MAPT* expression is mediated by long-range interactions with *cis*-regulatory elements

**DOI:** 10.1101/2023.03.07.531520

**Authors:** Brianne B. Rogers, Ashlyn G. Anderson, Shelby N. Lauzon, M. Natalie Davis, Rebecca M. Hauser, Sydney C. Roberts, Ivan Rodriguez-Nunez, Katie Trausch-Lowther, Erin A. Barinaga, Jared W. Taylor, Mark Mackiewicz, Brian S. Roberts, Sara J. Cooper, Lindsay F. Rizzardi, Richard M. Myers, J. Nicholas Cochran

**Affiliations:** HudsonAlpha Institute for Biotechnology, Huntsville, AL, USA; University of Alabama at Birmingham, Birmingham, AL, USA

**Keywords:** Alzheimer’s disease, *MAPT*, Gene regulation, Neuron, Enhancer

## Abstract

**Background:** Tauopathies are a group of neurodegenerative diseases driven by abnormal aggregates of tau, a microtubule associated protein encoded by the *MAPT* gene. *MAPT* expression is absent in neural progenitor cells (NPCs) and increases during differentiation. This temporally dynamic expression pattern suggests that *MAPT* expression is controlled by transcription factors and cis-regulatory elements specific to differentiated cell types. Given the relevance of *MAPT* expression to neurodegeneration pathogenesis, identification of such elements is relevant to understanding genetic risk factors.

**Methods:** We performed HiC, chromatin conformation capture (Capture-C), single-nucleus multiomics (RNA-seq+ATAC-seq), bulk ATAC-seq, and ChIP-seq for H3K27Ac and CTCF in NPCs and neurons differentiated from human iPSC cultures. We nominated candidate cis-regulatory elements (cCREs) for *MAPT* in human NPCs, differentiated neurons, and pure cultures of inhibitory and excitatory neurons. We then assayed these cCREs using luciferase assays and CRISPR interference (CRISPRi) experiments to measure their effects on *MAPT* expression. Finally, we integrated cCRE annotations into an analysis of genetic variation in AD cases and controls.

**Results:** Using orthogonal genomics approaches, we nominated 94 cCREs for *MAPT*, including the identification of cCREs specifically active in differentiated neurons. Eleven regions enhanced reporter gene transcription in luciferase assays. Using CRISPRi, 5 of the 94 regions tested were identified as necessary for *MAPT* expression as measured by RT-qPCR and RNA-seq. Rare and predicted damaging genetic variation in both nominated and confirmed CREs was depleted in AD cases relative to controls (OR = 0.40, p = 0.004), consistent with the hypothesis that variants that disrupt *MAPT* enhancer activity, and thereby reduce *MAPT* expression, may be protective against neurodegenerative disease.

**Conclusions:** We identified both proximal and distal regulatory elements for *MAPT* and confirmed the regulatory function for several regions, including three regions centromeric to *MAPT* beyond the well-described H1/H2 haplotype inversion breakpoint. This study provides compelling evidence for pursuing detailed knowledge of CREs for genes of interest to permit better understanding of disease risk.

## BACKGROUND

Aggregation of the microtubule associated protein tau (encoded by *MAPT*) is a defining pathological feature of neurodegenerative tauopathies like Alzheimer’s disease (AD) and tau-positive frontotemporal dementia (FTD) including progressive supranuclear palsy (PSP). Tau is highly abundant in the brain and functions to stabilize microtubules, which are critical for axonal growth and guidance.^1–3^ There are six isoforms of tau expressed in the brain and different isoforms have either three (3R) or four (4R) microtubule-binding domains that are regulated by phosphorylation at sites within or adjacent to the binding domains. In pathological conditions, tau undergoes hyperphosphorylation at these sites, permitting it to become unbound from microtubules and promoting tau oligomerization.^1, 4^ Hyperphosphorylation of tau leads to the mislocalization of tau from axons and into the soma and dendrites.^5, 6^ Somatodendritic mislocalization of tau occurs early in disease pathogenesis before neurodegeneration and mediates synaptic dysfunction leading to neuronal loss.^5–8^ Tau hyperphosphorylation–induced oligomers subsequently form large insoluble fibrils that are the major components of neurofibrillary tangles (NFTs). These intracellular NFTs can be found in the hippocampus during normal aging, but abnormal loads elsewhere in the cortex are pathological hallmarks of tauopathies.^1, 9, 10^ In animal models, reducing endogenous tau has been successful in preventing early mortality, cognitive deficits, and excitotoxicity even in the presence of amyloid-beta.^11–16^

While studies examining altered expression levels of *MAPT* in AD have produced mixed results ^17–20^, rare copy number events point to the relevance of *MAPT* gene expression in pathogenesis. For example, *MAPT* duplication resulting in 1.6- to 1.9-fold higher levels of *MAPT* mRNA leads to a tauopathy with AD-like presentation.^21^ Further, efforts to better understand genetic contributors to disease have resulted in large-scale genome-wide association studies (GWASs) which have identified *∼* 83 common variants that are associated with late-onset AD (LOAD).^22–26^ The majority of GWAS nominated variants are located in non-coding regions of the genome, pointing to the importance of understanding the roles these regions have in contributing to disease.^27, 28^ The leading hypothesis is that these regions are cis-regulatory elements (CREs), or enhancers, contributing to gene expression.^29, 30^ Large efforts, like ENCODE, have been made to identify CREs; however, experiments/data enabling CRE prediction in brain cell types are sparse, resulting in false negatives when attempting to identify enhancers for cell types specific to the brain. Chromatin looping, which can physically connect loci that are tens or hundreds of thousands of kilobases apart, also makes assigning non-coding GWAS single nucleotide variants (SNVs) to genes difficult. SNVs are typically assigned to the nearest gene in the locus, but because of chromatin looping enhancers may actually be in closer three-dimensional proximity with promoters of genes farther away in linear distance.^31–34^ A further complication is linkage disequilibrium (LD), in which multiple nearby alleles tend to be inherited together in haplotype blocks, which makes causal variant identification difficult. In fact, *MAPT* falls within one of the largest LD blocks known in the human genome, spanning 1.8 Mb.^35^ The locus can be divided into two haplotypes, defined by a 900 kb inversion event encompassing several genes, including *MAPT*. The “H1” haplotype, which is harbored by hg38, is the most common haplotype and has been associated with PSP, corticobasal degeneration (CBD), and Parkinson’s disease (PD).^36–41^ Additionally, the haplotype-tagging SNVs have been associated with AD.^22, 42^ In the H2 haplotype, which carries the inversion relative to hg38, the best evidence points to a reduction in *MAPT* expression and a lower risk for AD.^43^

Here, we nominate putative cis-regulatory elements of *MAPT* using orthogonal genomic approaches. We integrated single nucleus multiomics (snRNA-seq + snATAC-seq) from both cultured neurons and previously published dorsolateral prefrontal cortex (DLPFC) tissue^20^ to correlate chromatin accessibility with *MAPT* expression. We performed chromatin conformation assays (HiC and Capture-C) to determine the 3D interactions with the *MAPT* promoter in NPCs and matched differentiated neurons, as well as pure populations of human glutamatergic and GABAergic neurons. Finally, we manually inspected regions previously nominated by other studies,^44, 45^ and to nominate more proximal regions with 1) characteristic histone modifications (H3K27ac & H3K4me1), 2) high conservation, 3) transcription factor (TF) binding motifs, or 4) ENCODE cCRE annotation. We assessed these nominated regions for sufficiency to induce transcriptional activity through reporter assays and for necessity to *MAPT* expression using CRISPR interference (CRISPRi) experiments.

## METHODS

### Cell Lines

XCL4 Neural Progenitor Cells (NPCs) were obtained from StemCell Technologies. BC1 NPCs were obtained from MTI GlobalStem. All NPCs were maintained in NPC Media: 2:1 DMEM (high glucose, L-Glutamine, 100 mg/L Sodium Pyruvate): Ham’s F-12 Nutrient Mix, and supplemented with 50X serum-free B-27 Supplement (ThermoFisher Scientific). hFGF (40 *µ*g/mL), hEGF (20 *µ*g/mL), and heparin (5 ug/mL) were added daily to NPC media. iCell GlutaNeurons and iCell GABA Neurons were obtained from FujiFilm and maintained according to the manufacturer protocols for 14 days. HEK 293FT cells were obtained from ThermoFisher Scientific (R70007) and maintained in DMEM (high glucose, L-Glutamine, 100 mg/L Sodium Pyruvate) supplemented with 10% FBS, 1% Glutamax, 1% non-essential amino acids (NEAA), and 500 mg/mL Geneticin (G418 Sulfate) (ThermoFisher Scientific). KOLF2.1J and KOLF2.1J-hNGN2 iPSCs were obtained from the Jackson Laboratory and maintained in mTeSR Plus medium (StemCell Technologies). All cells were cultured at 37*^◦^*C with 5% CO2. KOLF2.1J NPCs were generated following the StemCell Technologies neural induction protocols.

### Neuron Differentiation

NPCs were differentiated according to the Bardy, et al. protocol.^46^ Cells were maintained in the neuronal differentiation medium for 14 or 21 days. iCell GlutaNeurons and GABAneurons were differentiated according to the manufacturer recommendations (FujiFilm Cellular Dynamics). Cells were maintained in their respective BrainPhys complete media for 14 days. KOLF2.1J-hNGN2 iPSCs were dissociated using Accutase and plated as single cells at 50,000 cells/cm^2^ in Induction Medium: DMEM/F12 medium with HEPES (ThermoFisher), 100x N2 supplement (StemCell Technologies), 100X Non-essential amino acids (NEAA, ThermoFisher), and 100X Glutamax (ThermoFisher). For plating cells, Induction Medium was supplemented with 5mM Y-27632 and 2 mg/mL Doxycycline. Induction medium supplemented with Doxycycline was renewed daily for two days. On day 3 neurites were present, and cells were renewed with Cortical Neuron Culture Media: BrainPhys neuronal medium (StemCell Technologies), 50X SM1 supplement (StemCell Technologies), 10 *µ*g/mL BDNF (StemCell Technologies), 10 *µ*g/mL NT-3 (StemCell Technologies), and 1 mg/mL Mouse Laminin (Gibco). On day 3, Cortical Neuron Culture Media was supplemented with 2 mg/mL Doxycycline. One-half media changes were performed every other day for a total of 14 days.

### Single nucleus multiomics

Nuclei from cultured neurons were isolated as detailed in 10x Genomics demonstrated protocol CG000375 Rev A with modifications. Cells were washed with cold 1X DPBS then 500 uL of NP-40 Lysis Buffer was added and cells were gently scraped off the dish while on ice, transferred to a 1.5 mL tube, and incubated 5 min on ice. Cells were centrifuged at 500 x g for 5 min at 4C and 500 uL 1X PBS + 1% BSA + DAPI + 0.5U/uL Protector RNAse Inhibitor (Roche) was added. Cells were spun again, gently resuspended 5X in 0.1X Lysis Buffer, and incubated 2 min on ice. After incubation, 1 mL of wash buffer was added and cells were immediately spun at 500 x g for 5 min at 4C. Nuclei were resuspended in <100 uL of Diluted Nuclei Buffer and counted using Countess FL II aiming for *∼* 3,000 - 5,000 nuclei/uL. Transposition, barcoding, and library preparation were performed according to the 10x Genomics Chromium Next GEM Single Cell Multiome protocol CG000338 Rev E. Protocols for single nucleus multiomics in DLPFC tissue were described in our previous publication.^20^

### Joint snRNA-seq and snATAC-seq workflow

Count matrices were obtained for each time point using cellranger-arc count (v2.0.1) then aggregated using cellranger-arc aggr. Low-quality cells were filtered on gene expression data (nFeatures > 200, nFeatures < 10,000, and mitochondrial percent < 5) and chromatin accessibility data (nucleosome signal < 2 and TSS enrichment > 2). Peaks that were present in less than 10 cells were removed from the ATAC matrix. RNA counts were normalized with SCTransform with mitochondrial percent per cell regressed out. Principal component analysis (PCA) was performed on RNA, and UMAP was run on the first 30 principal components (PCs). The optimum number of PCs was determined to be 30 PCs using an elbow plot. The ATAC counts were normalized with term-frequency inverse-document-frequency (TFIDF). Dimension reduction was performed with singular value decomposition (SVD) of the normalized ATAC matrix. The ATAC UMAP was created using the 2nd through the 50th LSI components. The weighted nearest neighbor (WNN) graph was determined with Seurat’s FindMultiModalNeighbors to represent a weighted combination of both modalities. The first 30 dimensions of the RNA PCA and the 2nd through the 50th dimensions from the ATAC LSI were used to create the graph. The WNN UMAP was created using the wknn (k=20). Clusters were identified from the wknn graph, and markers for each cluster were determined using a Wilcoxon rank-sum test. Clusters with marker genes that were enriched for ribosomal genes were filtered from the data (**Supplementary Tables 1-3**). After filtering low quality clusters, all normalization and dimension reduction were repeated as previously described.

### Differential Expression from snRNA-seq

Differentially expressed genes were determined for one cluster versus all other clusters using a Wilcoxon Rank Sum test. Only genes that were expressed in at least 10% of cells in a cluster were tested. Genes with a Bonferroni adjusted p-value < 0.01 were determined to be significant. Cluster markers were found for initial clusters to define and filter low quality clusters. Cluster DEGs were recalled with the same method after filtering.

### Gene Set Enrichment

The R package enrichR ^47–49^ was used for gene set enrichment analyses. Sets of positive DEGs for each cluster were used as input to look for enrichment in GO Biological Process 2021, GO Molecular Function 2021, GO Cellular Component 2021, and KEGG 2021 databases. Terms with an adjusted p-value less than 0.05 were considered to be enriched.

### Feature Linkage Analysis

Feature linkages were called using cellranger-arc. The maximum interaction distance was restricted to 1Mb, and feature linkages with a correlation score < 0.2 were removed and not used for downstream analysis. For feature linkage calculation, ATAC and GEX counts were normalized independently using depth-adaptive negative binomial normalization. To account for sparsity in the data, the normalized counts were smoothed by taking the weighted sum of the 30 closest neighbors from the KNN graph. The cell weights are determined by using a Guassian kernel transformation of the euclidean distance. Feature linkage scores were calculated by taking the Pearson correlation between the smoothed counts, while the significance of the correlation was determined using the Hotspot algorithm ^50^. For cultured multiomics, the ATAC peaks were called jointly across all time points. Cells from all time points were used to call links. For links called from AD and control tissue, ATAC peaks were called for each cell type and the union of these peaks was used to call links. Cells from all cell types were used in the feature linkage calculation.

### Chromatin Conformation Assays

HiC was performed in iCell GlutaNeurons, and Capture-C was performed in NPCs, Day 14 BrainPhys differentiated neurons, iCell GlutaNeurons and GABANeurons. All experiments were performed in triplicate with independent cell grow-ups of approximately 15 million cells for biological replication. For HiC, we followed manufacturer recommendations from the Arima HiC kit- User Guide for Mammalian cell lines (A510008, V.Oct.2019) using the Arima HiC Library Prep with Swift Biosciences Accel-NGS 2S plus DNA library kit (A510008, V. Nov2018). Libraries were sequenced using an Illumina Nova Seq S4 with XP kit for 2.5 billion reads total, which is *∼* 833 million reads per replicate for HiC.

Capture-C was performed in both iCell GlutaNeurons and GABANeurons following the HiC protocols, with the addition of the Agilent Technology SureSelect XT HS/SureSelect XT Low input target enrichment with pre-capture pooling protocol. 102 SureSelect DNA probes spanning a total of 2,073 bp (chr17:45892780-45893184 and chr17:45893527-45895196; hg38) were ordered from Agilent Technologies. The region between the two probe sets could not be synthesized because of low sequence complexity. Libraries were sequenced using an Illumina Nova Seq S4 with XP kit with *∼* 417 million reads per replicate.

NPC and Day 14 BrainPhys differentiated neuron Capture-C was performed using both the BC1 and XCL4 NPC lines with matched differentiated neurons. Capture-C was performed following the NG Capture-C Protocol (v2.4).^51^ For this experiment, enrichment for *MAPT* promoter contacts was performed by double capture using two biotinylated oligonucleotides targeting chr17:45892836–45892923 and chr17:45895092–45895179 (both hg38). Libraries were sequenced using an Illumina NovaSeq S4 with XP kit yielding *∼* 208 million reads per replicate.

### HiC and Capture-C Analysis

HiC data was analyzed using the Juicer pipeline (v1.6) with Juicer Tools (v1.22.01). Capture-C data was analyzed using the Juicer Tools pipeline (v1.21). Libraries from each replicate were first run individually. Restriction site positions were generated with *generate_site_positions.py* with Arima specified as the enzyme. Reads were aligned to hg38 for HiC, but only to chr17 of hg38 for capture-C. For both HiC and capture-C, all 3 replicates were merged to create a combined hic file with mega.sh. Loops were called on the Knight-Ruiz normalized combined hic files using HiCCUPs at a resolution of 5 kb. Window width and peak width were set to 20 and 10 respectively. Loops with a FDR < 0.2 were determined to be significant. Resulting loops were then filtered to a maximum interaction distance of 1Mb.

### Differential Capture-C Analysis

Knight-Ruitz normalized counts were pulled from the hic file for each replicate for Gluta and GABA capture-C at a resolution of 5kb and merged by contact positions. Counts that were not connected to the *MAPT* promoter were removed from the count matrix. Differential regions were tested using DESeq2 for 5kb bins. Neuron and NPC capture-C were processed using capC-MAP^52^ with the target set at the *MAPT* Promoter (chr17:45892837-45895177) and the restriction enzyme set as DpnII. Contact counts were taken from capC-MAP output at a step size of 500bp and window size of 1kb. Differential regions were tested using DESeq2 with cell line as a covariate. For both analyses, significant regions were defined as those with an adjusted p-value < 0.01. DESeq2 results were formatted as a bigwig for plotting using the directional p-value: -log_10_(adjusted p-value) x sign(log_2_FC).

### ATAC-seq Protocol

ATAC-seq was performed in the KOLF2.1J-hNGN2 cell line after 14 days of neuronal differentiation with two biological replicates per cell line. Our protocol is similar to those previously published by Buenrostro, et al.^53, 54^ Briefly, we harvested 100,000 cells by mechanical dissociation and washed with 50 *µ*L cold 1X PBS and centrifuged at 300xg for 5 minutes. The cell pellet was then resuspended in 50 *µ*L cold Lysis Buffer (10 mM Tris-HCl at pH 7.5, 10 mM NaCl, 3 mM MgCl2, 0.1% NP-40, 0.1% Tween-20, 0.01% Digitonin, and nuclease-free H2O) and incubated on ice for 3 minutes. After incubation, 1 mL of Wash Buffer (10 mM Tris-HCl at pH 7.5, 10 mM NaCl, 3 mM MgCl2, 0.1% Tween-20, and nuclease-free H2O) was added, samples were inverted gently to mix, and centrifuged at 500xg for 10 minutes at 4*^◦^*C. The supernatant was discarded and the remaining nuclei were resuspended in 50 *µ*L Transposition Mix (2X TD Buffer, TDE1, and nuclease-free H2O) and incubated at 37*^◦^*C on shaker at 1000 rpm for 30 minutes. Samples were immediately purified using the Qiagen MinElute Reaction Cleanup kit and eluted in a final volume of 11 *µ*L. Recovered DNA was then used to generate sequencing libraries using primers from the Nextera XT Index Kit (15055293) and Q5 Hot Start Master Mix and amplified (30 sec at 98*^◦^*C; [10 sec at 98*^◦^*C, 30 sec at 63*^◦^*C, and 72*^◦^*C for 1 min] ×10 cycles). Libraries were quantified with Qubit dsDNA HS Assay kit and visualized with BioAnalyzer High Sensitivity DNA Analysis kit (Agilent 5067-4626) and 2100 BioAnalyzer Instrument (Agilent). Libraries were sequenced using Illumina NovaSeq flowcell with 50-bp paired-end runs. Reads were processed using the standard ENCODE ATAC-seq pipeline (v1.7.0).

### ChIP-seq Protocol

ChIP-seq for H3K27Ac and CTCF were performed using chromatin from neurons differentiated from KOF2.1J NPCs (day 14) with biological replicates. Protocols for ChIP-seq are consistent with techniques previously described by our lab and available from the ENCODE Consortium (https://www.encodeproject.org/documents/73c95206-fc02-41ea-93e0-a929a6939aaf/).^55–57^ Antibodies targeting H3K27Ac (ActiveMotif, Cat: 39133) or CTCF (ActiveMotif, Cat:61311) were used. Libraries were prepared by blunting and ligating ChIP DNA fragments to sequencing adapters for amplification with barcoded primers (30 sec at 98*^◦^*C; [10 sec at 98*^◦^*C, 30 sec at 65*^◦^*C, 30 sec at 72*^◦^*C] x 15 cycles; 5 min at 72*^◦^*C). Libraries were quantified with Qubit dsDNA HS Assay kit and visualized with Standard Sensitivity NGS Fragment Analysis Kit (Advanced Analytical DNF-473) and Fragment Analyzer 5200 (Agilent). Libraries were sequenced using Illumina NovaSeq flow cell with 100bp single-end runs.

### ChIP-seq analysis

Prior to analysis, reads were processed to remove optical duplicates with clumpify (BBMap v38.20; https://sourceforge.net/projects/bbmap/) [dedupe=t optical=t dupedist=2500] and remove adapter reads with Cutadapt (v1.16)[-a AGATCGGAAGAGC-m 40]^8^. Input reads were capped at 40 million using Seqtk (v1.2; https://github.com/lh3/seqtk). Individual experiments were constructed following ENCODE guidelines (https://www.encodeproject.org/about/experiment-guidelines/) and analyzed with the chip-seq-pipeline2 processing pipeline (https://github.com/ENCODE-DCC/chip-seq-pipeline2). FASTQs were cropped to 50bp and all other parameters were run using the default setting. Final peaks were called using pseudoreps and the IDR naïve overlapping method with a threshold of 0.05.

### Plasmids

The pNL1.1.CMV [Nluc/CMV] and pGL4.23 [luc2/minP] vectors were obtained from Promega. Luciferase elements were generated by selecting 467 bp of the nominated region and both the forward and reverse complement sequences were ordered as gBlocks from Integrated DNA Technologies (IDT). Elements were cloned into the pGL4.23 [luc2/minP] vector digested with EcoRV by Gibson Assembly. Element insertion was confirmed by Sanger sequencing (MCLAB). Each element was individually prepped 3 times for a total of 6 individual plasmid preparations per nominated region. The pMD2.G, psPAX2, FUGW-H1-GFP-neomycin, and pLV hU6-sgRNA hUbC-dCas9-KRAB-T2A-Puro plasmids were obtained from Addgene (#12259, #12260, #37632, and #71236 respectively). sgRNA sequences were designed using https://benchling.com and ordered as premixed primer pools from IDT (**Supplementary Table 4**). The sgRNA sequences were then inserted into the pLV hU6-sgRNA hUbC-dCas9-KRAB-T2A-Puro plasmid by digesting with Esp3I and subsequent ligation.^59, 60^

### Lentivirus Production

HEK 293FT cells were plated at 70,000 cells/cm^2^ in poly-L-ornithine coated 6-well culture plates. The next day, the media was renewed with OptiMEM Reduced Serum Media supplemented with 300 mg D-glucose. Cells were transfected with 1 *µ*g pLV hU6-sgRNA hUbC-dCas9-KRAB-T2A-Puro plasmid with inserted sgRNA sequence using Lipofectamine LTX with Plus Reagent, following manufacturer recommendations. 48 hours post transfection, supernatant was harvested and filtered through a 0.45 *µ*m syringe filter into a 15 mL conical tube on ice.

### CRISPRi Experiments

NPCs were plated at 57,000 cells/cm^2^ in reduced growth factor Matrigel (Corning #354230) coated 12 well culture plates. The next day, culture media was renewed with NPC media supplemented with protamine sulfate (10 mg/mL). To transduce NPCs, 500 *µ*L of cold, filtered lentivirus was added to the NPC media and plates were centrifuged at 2000 rpm for 1 hour. 24 hours post transduction, cells were renewed with fresh NPC media with growth factors and 0.5 *µ*g/mL puromycin for selection of successfully transduced cells. Cells were selected for 24-48 hours, until the control well (transduced with FUGW-H1-GFP-neomycin) had minimal remaining living cells. NPCs were then swapped to neuronal differentiation media following the Bardy, et al. protocol^47^. Neurons were harvested for RNA isolation at day 14 of differentiation. sgRNAs were designed upstream of the *MAPT* TSS in the promoter region (*∼* chr17:45884000; hg38) as a positive control. sgRNAs targeting the AAVS1 safe harbor locus were used as non-targeting controls.

### RT-qPCR

*MAPT* expression was also determined by reverse transcription-quantitative polymerase chain reaction (RT-qPCR). cDNA was generated from 100 ng RNA using SuperScript IV Master Mix (Thermo 11756050). Two Taqman probes were used to measure *MAPT* relative abundance (Thermo Hs00213491_m1 and Hs00902194_m1). Taqman probes to AP1G1 (Thermo Hs00964419_m1) and GAPDH (Thermo Hs99999905_m1) were used as housekeeping controls (*AP1G1* was determined in pilot RNA-seq experiments to be a well expressed, low variability transcript between both NPCs and neurons). For each sample, Δ ΔCt values were calculated using the average of the median housekeeper Ct values. Samples with average housekeeper Ct values ± 1 standard deviation from the average of the other samples in each run were removed.

### 3’mRNA-seq Library Generation

RNA isolation was performed using the Norgen Total RNA isolation and RNase-Free DNase I kits (Norgen 17250 and 25720, respectively) and quantified using the Qubit RNA HS Assay Kit (Thermo Q32855). Libraries were prepared using the QuantSeq 3’ mRNA-Seq Library Prep Kit FWD for Illumina and UMI Second Strand Synthesis Module for QuantSeq FWD (Illumina, Read 1) from Lexogen (015.96 and 081.96, respectively). Libraries were quantified using the Qubit DNA HS Assay Kit (Thermo Q32854) and visualized with the BioAnalyzer High Sensitivity DNA Analysis kit (Agilent 5067-4626) and 2100 BioAnalyzer Instrument (Agilent). Libraries were sequenced by HudsonAlpha Discovery using Illumina NovaSeq S1 100 cycle or S4 200 cycle flowcells.

### RNAseq Analysis

UMIs were first extracted from the reads with UMI-tools extract with the extraction method set as regex. Reads were trimmed with bbduk then aligned to hg38 with STAR using the Lexogen recommended parameters for QuantSeq. Bams were deduplicated by UMI and mapping coordinate using UMI-tools dedup. Counts were generated with htseq-count using the intersection-nonempty method. The count matrix was normalized to counts per million (CPM). After normalization, each sample was scored based on the level of differentiation and astrocyte presence using Seurat’s *AddModuleScore*. Module scores were calculated using a list of markers for differentiation (*CACNA1C, ENO2, MAP2*) and astrocytes (*AQP4, SLC1A3*) (**Supplementary Fig. 1**). Differentially expressed genes (DEGs) were determined for each target region versus non-targeting controls for all genes in the *MAPT* locus (+1Mb). Differential expression was assessed with a linear model with differentiation score, astrocyte score, and batch as covariates. Genes with a p-value< 0.05 were determined to be significant.

### Nucleofection

iCell GlutaNeurons and KOLF2.1J-hNGN2 differentiated neurons were plated at 200,000 cells/cm^2^ in 24-well matrigel pre-coated plates. Cells were differentiated according to their respective protocols for 14 days. One hour before nucleofection, media was renewed to fresh media plus 500X Rock Inhibitor (Y27632). Neurons were nucleofected with 10 *µ* total DNA plasmid DNA using the AD1 4D-Nucleofector Y kit (Lonza V4YP-1A24) with the ED158 (iCell GlutaNeurons) and EH166 (KOLF2.1J-hNGN2 neurons) pulse settings. Per nucleofection, 9 *µ*g element cloned into pGl4.23 and 1 *µ*g pNL1.1.CMV [*Nluc*/CMV] were used. A transfection reaction of 9 *µ*g pGL4.23 [*luc2* /minP] and 1 *µ*g pNL1.1.CMV [*Nluc*/CMV] was used as a baseline control. Both vectors were also transfected as background controls (1 *µ*g) with pmaxGFP (9 *µ*g, Lonza). 24 hours post nucleofection, cells were renewed with fresh media without Rock inhibitor. Cell lysates were harvested by freezing at −80 *^◦^*C 48 hours post-transfection.

### Luciferase Assays

Luciferase assays were performed using the Nano-Glo Dual-Luciferase Reporter Assay System (Promega Cat#N1630) following the manufacturer’s protocol. Cell lysis was performed on the 24 well plate and aliquoted across 4 wells of a white 96-well plate for 4 technical replicates per biological replicate. Assays were completed in quadruplicate. Firefly luminescence was first normalized across the average plate luminescence and then normalized to the average control luminescence. For each biological replicate, the median fold luminescence value was determined for the four technical replicates. Four biological replicates were compared to the pGL4.23 [luc2/minP]/ pNL1.1.CMV [Nluc/CMV] control using an ordinary one-way ANOVA with Fisher’s LSD.

### Genetic Burden Analysis

Sequencing data were obtained from the 10th release of the Alzheimer’s Disease Sequencing Project. Data for chromosome 17 were filtered to the region of interest (hg38 chromosome 17 from 44.8 to 47.0 megabases, which is inclusive of *MAPT*, nearby genes, and all nominated regulatory regions) and annotated with dbSNP version 154, TOPMed Bravo freeze 8, CADD version 1.6, and the ENSEMBL GRCh38.99 gene model. Burden analysis was conducted on non-Hispanic white individuals only (the largest ancestral subset) to avoid confounding results from population stratification issues. PASS filter variants meeting criteria described in the results were aggregated, and individuals were counted as qualifying if they harbored 1 or more qualifying variants based on the filter conditions in the results (thus, if an individual harbored multiple qualifying variants, they were only counted once, though most individuals harbored only 1 qualifying variant). Differential analysis was conducted using a two-sided Fisher’s exact test comparing the number of cases and controls with and without a qualifying variant.

## RESULTS

### Identification of candidate CREs by single nucleus multiomics

To identify regions that may regulate *MAPT* expression, we used orthogonal approaches to nominate candidate genomic regions interacting with the *MAPT* promoter (**Fig. 1a**). First, we performed single-nucleus multiomics (snRNA-seq and snATAC-seq) using the 10x Genomics Multiome technology on nuclei isolated from neural progenitor cells (NPCs) and neurons differentiated from these NPCs for 14 and 21 days.^46^ This differentiation protocol produces a mixture of inhibitory and excitatory neurons and immature glial cells. Using this multiomics assay allows for direct mapping of gene expression and chromatin accessibility within the same nuclei without the need to computationally infer cell type identities computationally prior to cross-modality integration. We removed low quality nuclei and doublets (Methods), and we retained a total of 3,881 nuclei with an average of 1,293 per time point (range of 856 – 1,601). We detected a median of 3,559 genes and 55,996 ATAC fragments per cell. We performed normalization and dimensionality reduction for snRNA-seq and snATAC-seq data using Seurat (v4) and Signac (v5), respectively. We used weighted-nearest neighbor (WNN) analysis to determine a joint representation of expression and accessibility and identified 3 distinct clusters (**Fig. 1B, Supplementary Fig. 2a-d**). Cluster 1 represents a mostly (98.1%) NPC cluster and is defined by expression of *SOX5*, which regulates embryonic development and cell fate determination.^61^ Cluster 2 is a mixture of neurons differentiated for 14 (43.8%) and 21 days (56.2%) and is defined by expression of *NEAT1*, which in iPSC-derived neurons has been shown to directly regulate neuronal excitability.^62^ Cluster 3 is also a mixture of time points of differentiated neurons (19.3% Day 14 and 80.7% Day 21) and is defined by expression of *DCX*, which encodes doublecortin that is expressed in migrating neurons through the central and peripheral nervous system during embryonic and postnatal development, representing an immature neuron population.^63^ Cluster 3 also is the only cluster that expresses *MAPT*. This clustering was expected given the resulting mixed culture produced from this differentiation protocol. Differentially expressed genes (DEGs) were identified for each cluster by comparing to the other two remaining clusters (Supplementary Table 3). A total of 792 DEGs were identified, including *MAPT* (log_2_(Fold Change) = 0.75; adjusted p =5.39×10^-124^).

**Figure 1.**
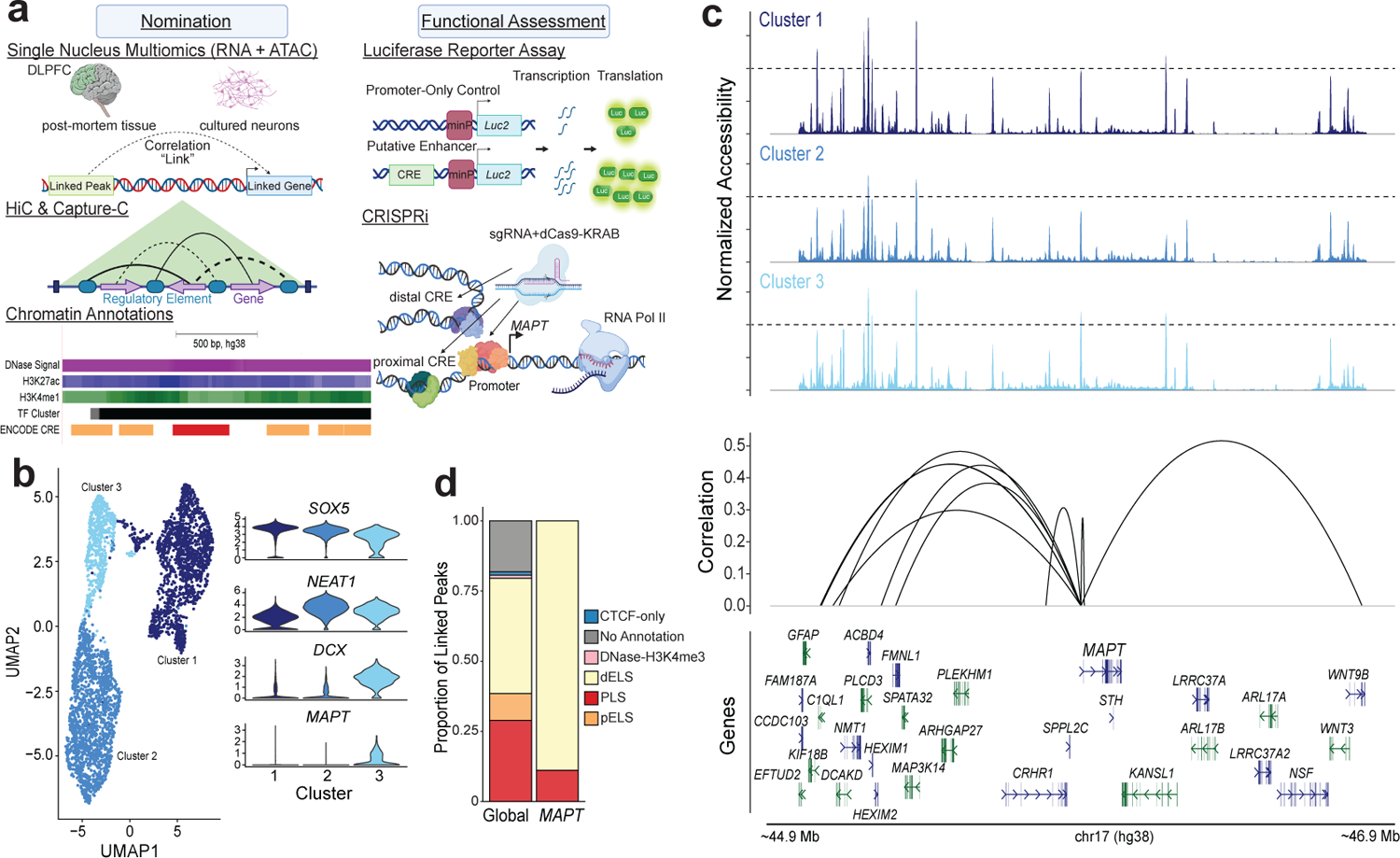
Nomination of candidate CREs by snRNA- & snATAC-seq in iPSC-derived neuronal cultures. **a)** Experimental design. **b)** Left:UMAP visualization of the weighted nearest neighbor (WNN) clustering of single nuclei colored by cluster assignment. Right: Violin plot of expression of key genes by cluster assignment **c)** Linkage plot for all links to *MAPT*. Top panels: coverage plot of normalized accessibility separated by cluster assignment (light blue = NPC; blue = intermediate differentiation; dark blue = immature neurons). Bottom panel: significant peak-gene links. Arc height represents the strength of correlation shown as absolute correlation values **d)** Comparison of proportion of linked peaks by ENCODE regulatory element annotation of global and *MAPT* links.

Using the cellranger-arc (v2.0) analysis pipeline, we measured correlations, or “links”, between gene expression and chromatin accessibility to nominate CREs. A “linked peak” is an ATAC-seq peak whose accessibility across all nuclei is significantly correlated with the expression of a “linked gene” (**Fig. 1a, top panel**). We restricted this correlation analysis to consider only peaks within 1 megabase (Mb) of each transcription start site (TSS), given previous studies that the vast majority of distal regulatory elements are less than 1 Mb from their target genes (see Methods).^64–72^ A total of 54,879 links were found, including 9 *MAPT*-specific links (**Fig. 1c, Table 1 and Supplementary Table 5, 6**). We furthermore overlapped the linked peaks with ENCODE curated CREs.73 The linked peaks were enriched for promoter-like sequences (Odds Ratio (OR) = 6.11; p < 2.2×10-16) and proximal enhancer-like sequences (OR = 1.59; p < 2.2×10-16), with the majority (8 of 9, 88.9%) of *MAPT* linked peaks overlapping distal enhancer-like sequences (**Fig.1d**). ATAC peaks can be linked to one or more genes; we identified a median of 1 link per peak globally (range 1 – 31) and 2 links per peak for *MAPT* linked peaks (range 1 – 3; **Supplementary Fig. 2e**). Additionally, the mean peak distance from gene TSS was *∼* 440 kb globally, but for *MAPT* linked peaks, the peaks tended to be farther from the TSS, with an average distance of *∼* 670 kb (Supplementary Fig. 2f).

**Table 1:**
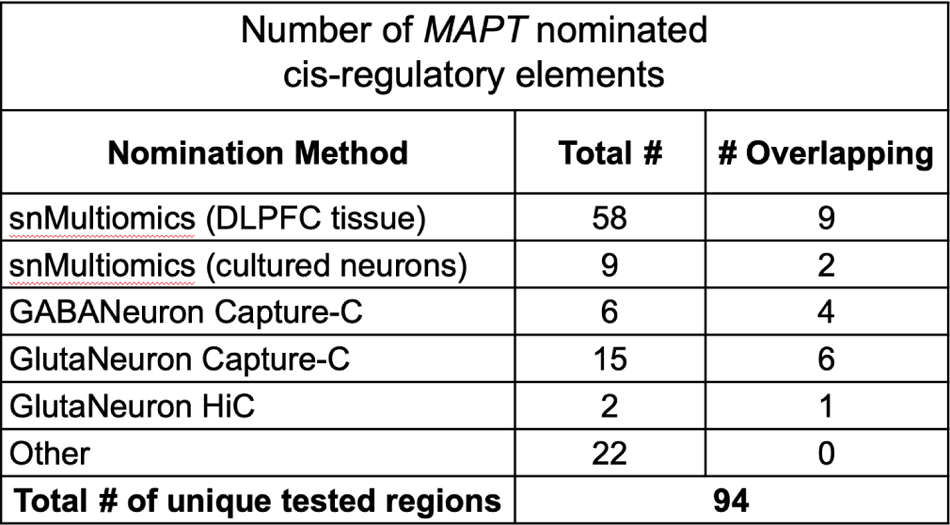
Total number of MAPT nominated cis-regulatory elements by nomination method. For each nomination method, the total number of nominated regions and number of regions that were also nominated by another method are shown. The total number of unique regions tested across methods is shown at the bottom.

In addition to the single nucleus multiomics data generated from cultured neurons for this study, we also identified *MAPT* links from our previously published data. ^20^ Using nuclei from DLPFC of 7 AD and 8 control donors, we identified cell type- and disease-specific CREs and their target genes. Focusing on the *MAPT* locus, we expanded the original restriction of 500 kb from each gene’s TSS in our previous study to 1 Mb to match the current analysis. We identified 58 *MAPT* links; 28 of these links were called in both AD and controls (common), while there were 26 AD-specific and 4 control-specific links (**Supplementary Fig. 3a and Supplementary Table 7, 8**). We found that many of the linked peaks were negatively correlated with *MAPT* expression. As with the published analysis method, links were called across all cells in the dataset. These negative correlations are likely driven in part by high accessibility of cCREs in cell types where *MAPT* is not expressed and the low variability of *MAPT* expression in neurons (example shown in **Supplementary Fig. 3b**). However, we included all correlated regions in the validation set to comprehensively evaluate all potential links. We overlapped the *MAPT* linked peaks with ENCODE CREs and found that they were significantly enriched for both proximal (OR = 4.50; p = 1.58×10-13) and distal enhancer-like sequences (OR = 1.98; p = 0.011) and promoter-like sequences (OR = 4.51; p = 1.58×10-13; **Supplementary Fig. 3c**).^73^ *MAPT* linked peaks had a median of 7.5 (range 1 – 15) links, while globally the median links per peak was only 3 (range 1 – 31; **Supplementary Fig. 3d**). *MAPT* links also tended to be farther away from the TSS in this data set with an average peak distance of *∼* 500 kb compared to *∼* 440 kb globally (**Supplementary Fig. 3e**).

### Candidate CREs identification by structural analysis and chromatin marks

Enhancers can exert their function by being in physical contact with the target gene’s promoter, despite large intervening sequencing distances, via chromatin looping.^32, 34, 64, 74^ We sought to identify looping events involving the *MAPT* promoter using HiC and Capture-C assays. We performed HiC in iCell GlutaNeurons, which produce a > 90% pure population of human glutamatergic (excitatory) neurons. Using the Juicer Tools pipeline (v1.21) and HiCCUPs, we identified 15,918 loops genome-wide with 2 of these loops contacting the *MAPT* promoter region (chr17: 45830001 – 45835000 [hg38]; **Fig. 2a and Supplementary Fig. 4a; Table 1 and Supplementary Table 9**). When overlapping loop ends with ENCODE annotations, about half are interactions with enhancer-like sequences (enhancer:enhancer 25.2% and enhancer:promoter 23.8%) and 17.7% are promoter:promoter interactions (**Fig. 2b and Supplementary 4b**).^73^ Both loops interacting with the *MAPT* promoter are regions annotated as enhancer-like sequences.

**Figure 2.**
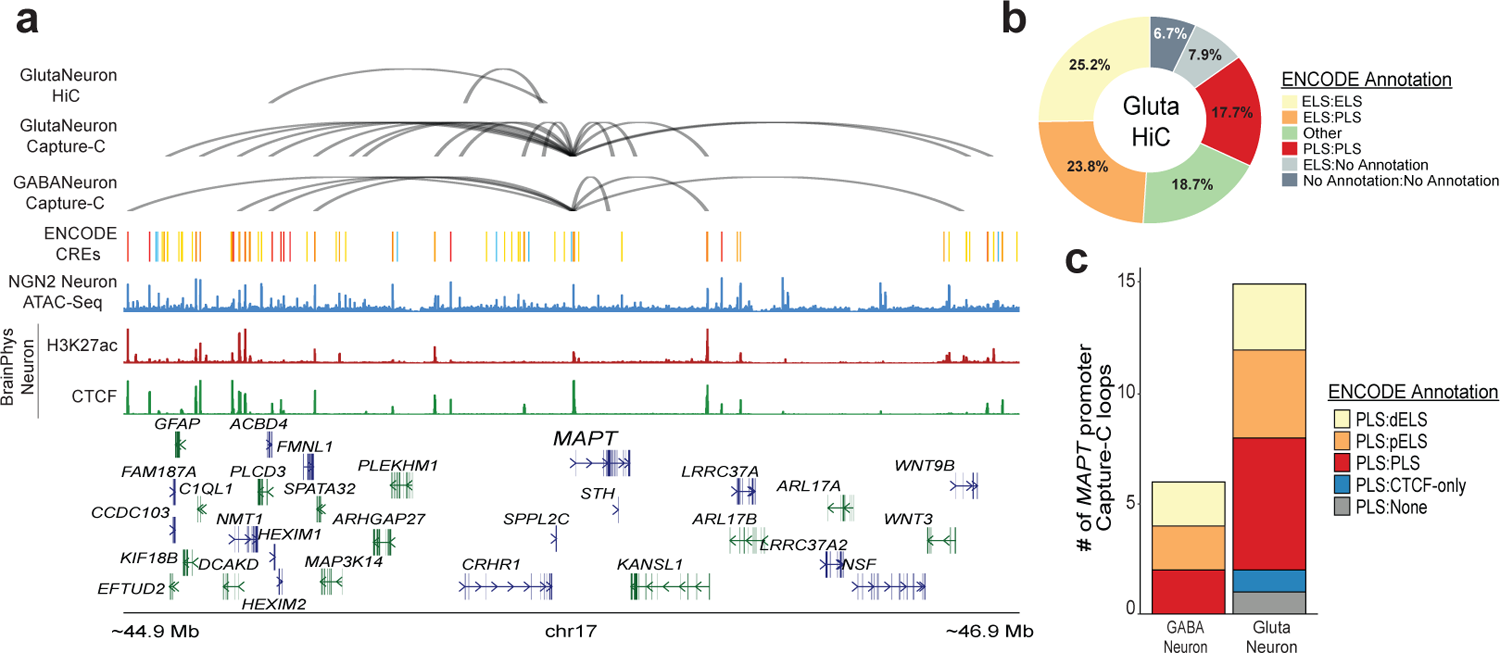
Nomination of candidate CREs using HiC and Capture C in pure excitatory and inhibitory neuron cultures. **a)** Loop plot of HiCCUPs nominated loops to the *MAPT* promoter. Top panel: Nominated loops by dataset. Top: HiC loops to the *MAPT* promoter from iCell GlutaNeuron cultures. Middle: Capture-C loops to the *MAPT* promoter from iCell GlutaNeuron cultures. Bottom: Capture-C loops to the *MAPT* promoter from iCell GABANeuron cultures. Bottom panel: Top: Track of ENCODE annotated candidate CREs. Middle: ATAC-seq peaks from KOLF2.1J-hNGN2 differentiated neurons. Bottom: H3K27ac and CTCF ChIP-seq peaks from Day 14 BrainPhys differentiated neurons. **b)** Proportion of HiC global loop calls by ENCODE annotation. **c)** Proportion of Capture-C *MAPT* promoter loops by ENCODE annotation for both iCell GlutaNeurons and GABANeurons.

To more sensitively detect regions specifically interacting with the *MAPT* promoter, we used AgilentTechnologies SureSelect DNA probes spanning a 2kb region (chr17: 45892780 – 45893184 and chr17: 45893527 – 45895196; hg38) to perform Capture-C in iCell GlutaNeurons and iCell GABANeurons (> 95% pure population of GABAergic (inhibitory) neurons) cultures. Loops were called on the Knight-Ruiz normalized combined hic file using HiCCUPs at a resolution of 5kb with the *MAPT* promoter region defined as chr17: 45890001 – 45895000. Resulting loops were then filtered to a maximum interaction distance of 1Mb, matching the chosen restriction space for single nucleus multiomics nominations. We identified 15 and 6 *MAPT* promoter contact loops in GlutaNeuron and GABANeurons, respectively (**Fig. 2a and Supplementary Fig. 4a; Tables 1, SupplementaryTables 10, 1**). About half of these loops are regions annotated as either proximal or distal enhancer-like sequences with the majority of the remaining loops being promoter-like sequences (**Fig. 2c**).

We also nominated candidate CREs by manual inspection using the UCSC genome browser (http://genome.ucsc.edu) incorporating ENCODE chromatin annotations (DNase Hypersensitivity, H3K27ac, and H3K4me1)^75^, transcription factor clusters (ENCODE)^75, 76^, conservation (Vertebrate Multiz Alignment and Conservation [100 Species]), and previously published data.^44, 45^ Limiting the search area to the 1 Mb upstream or downstream of the *MAPT* promoter, we nominated an additional 22 regions not overlapping any of the experimental nomination methods (“Other” Method **Table 1, Supplementary Table 12**). Of these nominated regions, most (56%) overlap distal enhancer-like sequences, but many are not annotated by ENCODE. However, since ENCODE annotations are defined by non-neuronal cell types, these nominated regions may be CREs specific to neuronal cell types (**Supplementary Fig. 4c**).

### Differential analysis of CREs nominated by structural analysis

In the chosen culture system, *MAPT* is very lowly expressed at the NPC stage, but is more highly expressed in the differentiated neuron state (**Fig.3a**). This expression switch makes this model very useful in examining *MAPT* regulation. To interrogate cCREs controlling this biological switch in *MAPT* expression, we performed differential analysis of Capture-C data generated in NPCs and differentiated neurons. For differential Capture-C, we chose bins of 500 bp, and we identified 49 regions differentially interacting with the *MAPT* promoter between NPC and neurons (**Supplementary Table 13**). Of these interacting regions, 27 were specific to neurons (**Fig. 3b**).

**Figure 3.**
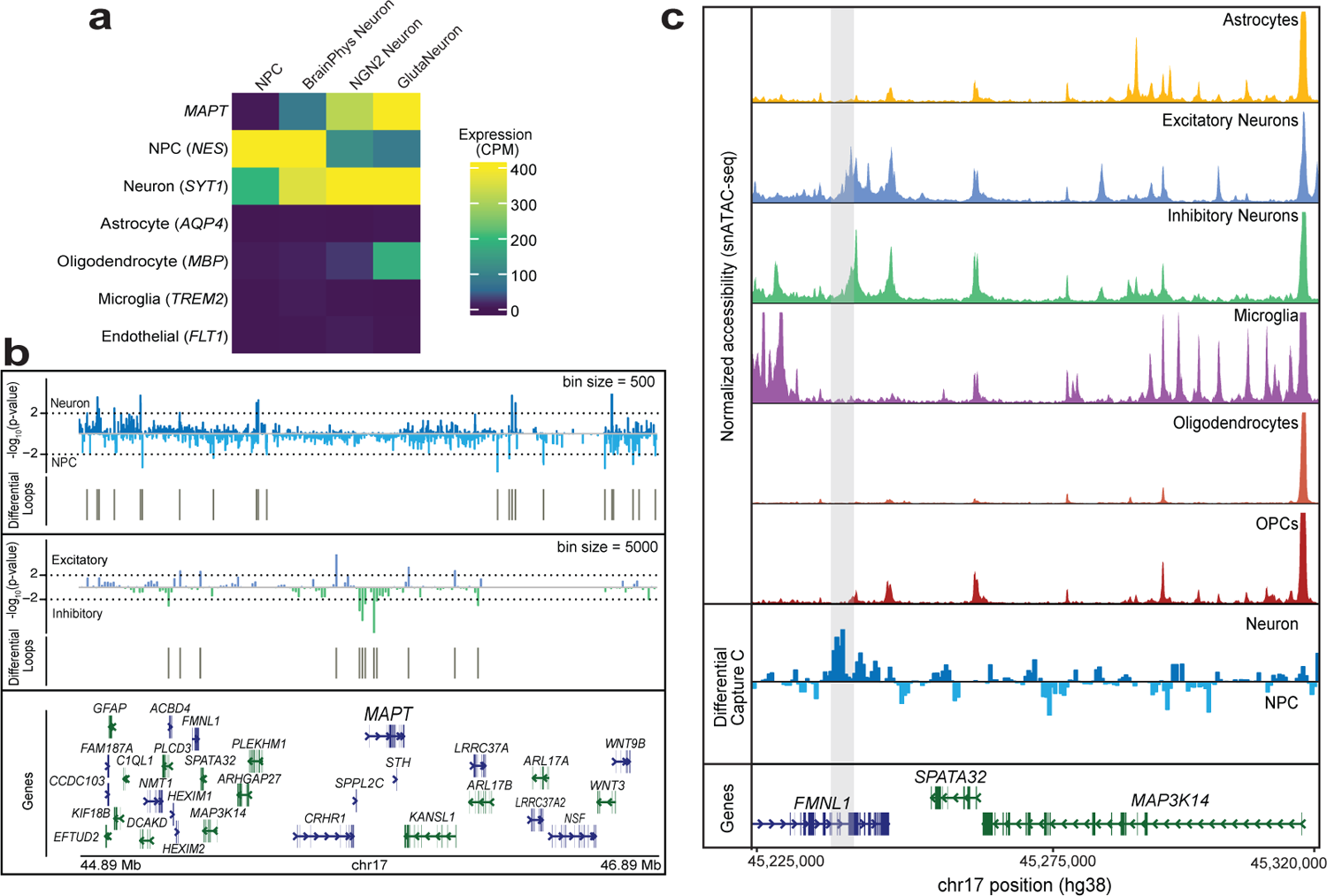
Identification of neuron-specific candidate regulatory regions. **a)** Heatmap showing expression of *MAPT* and cell type marker genes in NPCs, Day 14 BrainPhys differentiated neurons (BrainPhys Neurons), KOLF2.1J-hNGN2 Neurons (NGN2 Neuron), and iCell GlutaNeurons. **b**) Differential Capture-C directional p-value track. Top panel: Comparison of NPCs and Day 14 BrainPhys Neurons. Middle panel: Comparison of iCell GlutaNeurons (Excitatory) and iCell GABANeurons (Inhibitory). **c)** Zoom in of chr17: 45,225,000-45,320,000. Top panels: Chromatin accessibility across cell types generated by snATAC-seq in DLPFC nuclei. Bottom panels: Differential capture-C comparing BrainPhys differentiated neurons and NPCs with the neuron-specific region highlighted in gray.

One of these neuron-specific regions falls within a single nucleus multiomics link located 933,730 bp upstream of *MAPT* in the 3’ UTR of *C1QL1*. We also performed differential analysis of the interacting regions identified in excitatory and inhibitory neurons and found that 12 regions differentially interacted with the *MAPT* promoter (**Fig. 3b and Supplementary Table 14**). Of these differential regions, nine overlapped the nominated *MAPT* regulatory regions with five being nominated from single nucleus links. For the remaining overlapping regions, three were previously identified in the excitatory neuron Capture-C and one was identified in inhibitory neuron Capture-C.

One neuron-specific interacting region identified was located >650 kb upstream of the *MAPT* promoter in the *FMNL1* gene body (**Fig. 3c**). We evaluated chromatin accessibility within this locus using snATAC-seq data generated previously.^20^ We found that this region was specifically accessible in both excitatory and inhibitory neurons from adult DLPFC tissue, providing further evidence that this region likely functions as a neuron-specific regulatory element. In agreement with our findings, a previous study also found that this region harbored putative neuron-specific regulatory elements.^45^

### Functional assessment of candidate CREs

We sought to functionally assess the 94 nominated regions using human neurons differentiated from NPCs (BrainPhys), glutamatergic neurons generated from KOLF2.1J-hNGN2 iPSCs, and iCell GlutaNeurons. All of these models highly express the neuronal marker gene SYT177 and *MAPT* (**Fig. 3a**). We first tested 39 of the nominated regions for enhancer-like activity by testing for sufficiency to induce transcriptional activity using a luciferase reporter assay (**Supplementary Table 15**). We performed reporter assays in pure cultures of human iPSC-derived glutamatergic (excitatory) neurons (iCell GlutaNeurons and KOLF2.1J-hNGN2).^78, 79^ We selected regions with high levels of conservation, high DNase hypersensitivity signal, histone modifications characteristic of active elements (H3K27ac and H3K4me1), and overlapping transcription factor motif clusters^75^ and ENCODE candidate CREs.^73^ We excluded regions annotated as promoter-like sequences because of the prior hypothesis that these regions would be active in a luciferase assay (as expected for a promoter sequence) and therefore would not provide useful information on the sufficiency of these sequences to induce transcription. Of the nominated regions, 21 were tested in iCell GlutaNeurons. During the course of this study, the KOLF2.1J-hNGN2 cell line was made publicly accessible and produced pure excitatory neurons.^78, 79^ These neurons were more experimentally tractable than iCell GlutaNeurons, so the remaining 18 were tested in KOLF2.1J-hNGN2 derived neurons (**Supplementary Fig. 5a,b**). Eleven of the regions significantly increased activity of the luciferase reporter (**Fig. 4a and 4b**).

**Figure 4.**
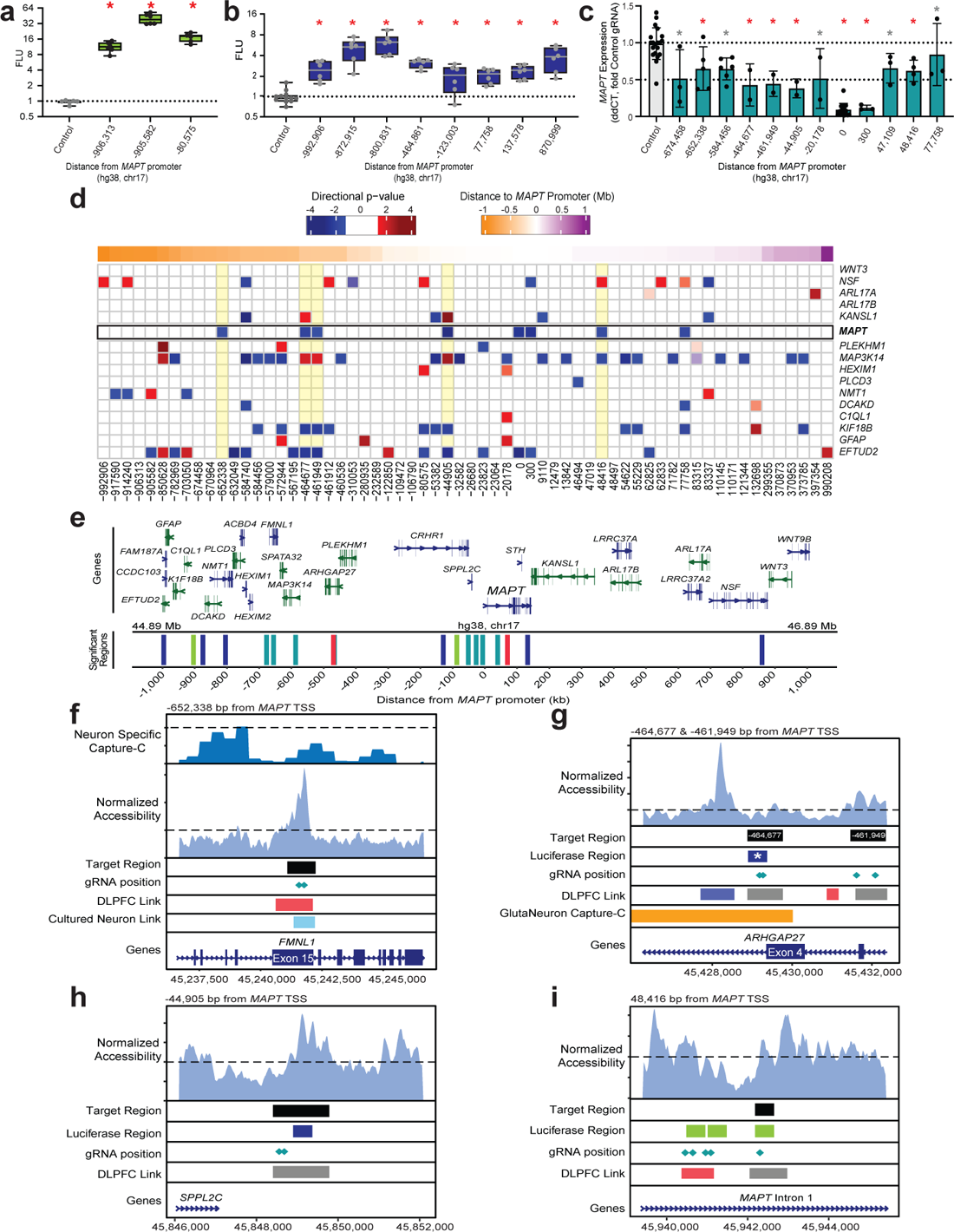
Functional validation of nominated CREs. **a.** Boxplots showing statistically significant (*p<0.05, ANOVA with Fisher’s LSD) elements tested in luciferase assays in iCell GlutaNeurons. **b.** Boxplots showing statistically significant (*p<0.05, ANOVA with Fisher’s LSD) elements tested in luciferase assays in KOLF2.1J-hNGN2 neurons. **c.** RT-qPCR of MAPT expression. Barplots show statistically significant regions tested in Day 14 BrainPhys differentiated neurons by CRISPRi experiments. Red asterisk indicates regions significant by 3’ mRNA-seq and RT-qPCR. Gray asterisk indicates regions significant in either RT-qPCR or 3’ mRNA-seq. (*p<0.05, ANOVA with Fisher’s LSD).**d.** Heatmap showing differential expression of MAPT and all genes expressed (cpm > 30) within 1 Mb of MAPT TSS. Left y-axis indicates the distance of the midpoint of the target region tested in CRISPRi experiments from the MAPT promoter (chr17:45894000). **e.** Browser view of significant nominated regions. Green: iCell GlutaNeuron luciferase assay. Blue: KOLF2.1J-hNGN2 neuron luciferase assay. Teal: CRISPRi. Red: significant in luciferase and CRISPRi. **f. – i.**Key CREs regulating MAPT expression. Zoom in on regions validated by qPCR and CRISPRi (**f.** −652,338, **g.** −464,677 & −461,949, **h.** −44,905, **i.** 48,416). Top panel: Normalized Accessibility from KOLF2.1J-hNGN2 differentiated Neurons. Bottom panel: Tracks of design region, luciferase region (Green: tested in iCell Gluta Neurons; Blue: tested in KOLF2.1J-hNGN2 neurons; *p<0.05, ANOVA with Fisher’s LSD), gRNA position, and nominated regions (DLPFC Links: Blue: control-specific link; gray: common link; Red: AD-specific link).

Since we tested a subset of the nominated regions for luciferase activity, we also assessed publicly accessible Massively Parallel Reporter Assay (MPRA) data to determine if the nominated regions had enhancer-like activity. Seventeen of the nominated regions overlap significant elements from either the Cooper, et al.^44^ MPRA or the van Arensbergen, et al.^80^ SuRE-seq datasets. Given that these datasets are assessing cCREs across the genome and in non-neuronal cells, it is not unexpected that the overlap with our regions is only *∼* 18% as genome-wide assays may have lower sensitivity than targeted assays for *MAPT*. Therefore, we also overlapped nominations with HiChIP for H3K27ac data generated from multiple brain regions by Corces, et al.^45^ Eighteen of the nominated regions overlapped a H3K27ac HiChIP peak (q < 0.01), including 5 regions overlapping either of the two MPRAs (**Supplementary Table 16**). Importantly, the region corresponding to the element 77,758 bp away from the *MAPT* promoter that had significant luciferase activity in KOLF2.1J-hNGN2 neurons was also significant in both MPRAs analyzed and overlapped a H3K27ac HiChIP peak (**Fig.4b**). Together these pieces of evidence support that this region has enhancer-like activity and interacts with the *MAPT* promoter.

We used CRISPRi to determine the target gene of the nominated regulatory regions, as well as their necessity for that gene’s expression. CRISPRi employs a catalytically inactive Cas9 enzyme (dCas9) fused to a Krüppel-associated box (KRAB) domain that recruits transcriptional repressors, and it has been previously shown to be effective at reducing gene expression in neurons with minimal off-target effects.^81^ At least one guide RNA (gRNA) was tested per candidate region, and positive controls were designed to target a region encompassing the *MAPT* promoter and TSS. We also designed gRNAs to both GFP, which is not expressed in our cell lines, and the AAVS1 safe harbor locus as negative controls where gRNAs should have no effect on gene expression (the AAVS1 safe harbor locus is on a different chromosome [19 vs. *MAPT* on 17] and has been extensively characterized to have limited cellular effects when edited).^82–84^ gRNAs were introduced into NPCs that were subsequently differentiated into neurons and harvested after two weeks. We isolated RNA and performed both quantitative reverse-transcription PCR (qRT-PCR) and 3’ mRNA-sequencing to assess gene expression changes for 62 of the nominated regions (**Supplementary Fig. 5c, Supplementary Table 17**). Introduction of the gRNAs for the remaining regions either resulted in cell death or RNA-seq data did not meet quality control metrics. We observed robust knockdown of *MAPT* expression when targeting the promoter and 300 bp downstream. We observed significant knockdown of *MAPT* after repressing 11 other regions, six of which were significant in either RNA-seq or RT-qPCR, but not both (**Fig. 4c, gray asterisks, and Fig. 4d**). The remaining five regions were confirmed in both 3’ mRNA-sequencing and RT-qPCR and are thus identified as regulators of *MAPT* (**Fig. 4c-e**).

The validated regulatory regions span from >650 kb upstream to *∼* 50 kb downstream of the *MAPT* promoter, and three of these are beyond the centromeric H1/H2 inversion breakpoint. The gRNA target region located 652,338 bp upstream of the *MAPT* promoter resulted in *∼* 35% knockdown of *MAPT* expression (**Fig. 4c, f**). This region was nominated in both cultured neuron and DLPFC single nucleus multiomics datasets; it lies within exon 15 of *FMNL1*, although *FMNL1* is not expressed in our cell lines (average cpm < 1). Two regions within the *ARHGAP27* locus each resulted in more than 50% knockdown of *MAPT* expression (**Fig. 4c**). Targeting both of these regions also significantly reduced expression of *EFTUD2* (log_2_FC = −0.414; p = 0.004, log_2_FC = −0.515; p =0.003) and *KIF18B* (log_2_FC = −1.367; p = 0.006, log_2_FC = −1.411; p = 0.006), while *MAP3K14* (log_2_FC = 0.766; p = 0.011, log_2_FC = 1.114; p = 0.015) expression was significantly increased (**Fig.4d,Supplementary Table 11**). *ARHGAP27* is very lowly expressed in the chosen culture system (average cpm *∼* 2), so we could not evaluate any effect on its expression. These regions were both nominated by DLPFC single nucleus multiomics, and the linked peaks identified were also linked to MAP3K14. The region 464,677 bp upstream from the *MAPT* TSS was additionally nominated by Capture-C and showed cell type-specific contact with the *MAPT* promoter in excitatory neurons. This region also had enhancer-like activity in the KOLF2.1J-hNGN2 neuron luciferase dataset (**Fig. 4b[Region −464,861], g**). Together these data indicate that this region is an enhancer regulating *MAPT*. The closest validated region (Region −44,905) lies within the first intron of *MAPT-AS1* and was nominated by DLPFC single cell multiomics with links to 11 genes including *MAPT*, *MAPT-AS1*, and *KANSL1* (**Fig. 4h**). Targeting this region resulted in > 60% reduction of *MAPT* expression, significant reduction in *EFTUD2* (log_2_FC = −0.479; p = 0.0002) and *KIF18B* (log_2_FC = −0.932; p = 0.008), and significant increase in *MAP3K14* (log_2_FC = 0.934; p = 0.0025) and *KANSL1* (log_2_FC = 1.135; p = 4.74 x 10-5) expression (**Fig. 4c,d**). The only downstream region validated in both RNA-seq and RT-qPCR was 48,416 downstream of the *MAPT* promoter, in the first intron of *MAPT* **(Fig. 4i**), a region with established importance due to the presence of the H1c tagging variant rs242557.^85^ This region was nominated by single nucleus multiomics and is linked to 10 genes, including *MAPT* and *NSF*. Targeting this region resulted in *∼* 40% reduction of *MAPT* expression, significant reduction in *MAP3K14* (log_2_FC = −0.82; p = 0.0169), and significant increase in *NSF* (log_2_FC = 0.232; p = 0.0351) expression (**Fig. 4c,d**). This region was previously established as interacting with *MAPT* based on its overlap with a H3K27Ac HiChIP peak.^45^

### MAPT cCREs are depleted of rare, deleterious variants in ADSP

We next asked if the regions we identified exhibited differential burden of rare genetic variation between AD cases and controls. We evaluated individuals in the Alzheimer’s Disease Sequencing Project case/control dataset with filtering conditions described in **Table 2** for all candidate regulatory regions. Very rare (allele frequency less than 1 in 100,000 in the TOPMed Bravo population database as well as an allele count of 1 in the Alzheimer’s Disease Sequencing Project (ADSP) set of 33,532 individuals) and predicted damaging (CADD>20) variants were depleted in cases vs. controls across implicated regulatory regions for *MAPT* (OR = 0.40, p = 0.004; qualifying variants are described in **Supplemental Table 18**). This effect direction is expected, as we would hypothesize that rare and damaging genetic variation in these regulatory elements would be associated with a reduction of their function, which in turn would lead to lower expression of *MAPT* and therefore be protective against neurodegenerative disease. Importantly, it is clear that experimental identification of regulatory regions implicated for *MAPT* adds information, as simply assessing the burden of genetic variation across the entire region using the same filtering conditions does not reveal a depletion of burden of qualifying genetic variants in cases.

**Table 2:** Burden analysis of rare genetic variants in ADSP data from non-Hispanic white individuals. Rare genetic variation at the hg38 chr17:44.8–47.0 Mb region spanning *MAPT*, nearby genes, and associated regulatory elements in Alzheimer’s Disease Sequencing Project (ADSP) data from self-reported non-Hispanic white individuals. Very rare (allele frequency less than 1 in 100,000 in the TOPMed Bravo population database as well as an allele count of 1 in the ADSP set of 33,532 individuals) and predicted damaging (CADD>20) variants are depleted in cases vs. controls across implicated regulatory regions for *MAPT*, but not across the region as a whole without enrichment for putative regulatory elements (last line of table). p value is indicative of a two-sided Fisher’s exact test. CADD: scaled Combined Annotation Dependent Depletion score. AF: allele frequency. OR (95% CI): odds ratio (95% confidence interval). Bonf.: Bonferroni-adjusted. kb: kilobase. KD: knockdown. LoF: loss-of-function. Mb: megabase.

| Segmentation | CADD<br>> | TOPMed AF<br>Threshold | Cases<br>w/ | Cases<br>w/o | Ctrl<br>w/ | Ctrl<br>w/o | OR<br>(95% CI) | p | Bonf.<br>p |
| --- | --- | --- | --- | --- | --- | --- | --- | --- | --- |
| All candidate regions<br>(83 kb total) | 10 | 1 in 1,000 | 985 | 4628 | 679 | 3201 | 1.00 (0.90–1.12) | 0.956 | 1.00 |
| All candidate regions<br>(83 kb total) | 10 | 1 in 10,000 | 534 | 5079 | 346 | 3534 | 1.07 (0.93–1.24) | 0.331 | 1.00 |
| All candidate regions<br>(83 kb total) | 10 | 1 in 100,000 | 189 | 5424 | 137 | 3743 | 0.95 (0.76–1.20) | 0.688 | 1.00 |
| All candidate regions<br>(83 kb total) | 20 | 1 in 1,000 | 87 | 5526 | 77 | 3803 | 0.78 (0.56–1.07) | 0.128 | 0.77 |
| All candidate regions<br>(83 kb total) | 20 | 1 in 10,000 | 49 | 5564 | 50 | 3830 | 0.67 (0.44–1.02) | 0.051 | 0.31 |
| All candidate regions<br>(83 kb total) | 20 | 1 in 100,000 | 15 | 5598 | 26 | 3854 | 0.40 (0.20–0.78) | 0.004* | 0.02* |
| hg38 chr17:44.8–47.0<br>Mb region | 20 | 1 in 100,000 | 208 | 5405 | 153 | 3727 | 0.94 (0.75–1.17) | 0.549 | NA |
| RNA-seq & qPCR KD<br>(6 kb total) | 20 | 1 in 100,000 | 1 | 5612 | 5 | 3875 | 0.14 (0.00–1.24) | 0.045* | NA |
| *MAPT* canonical<br>coding LoF | 10 | 1 in 1,000 | 3 | 5610 | 3 | 3877 | 0.69 (0.09–5.16) | 0.694 | NA |

Given the observation of depletion of very rare and predicted damaging genetic variation in cases across all nominated regions, we were particularly interested in the regions that showed clear knockdown when targeted with CRISPR/dCas9-KRAB. While numbers were small for qualifying variation in these elements (1 case and 5 controls), the depletion in cases persisted with nominal significance (OR = 0.14, p = 0.045).

Finally, under the hypothesis that rare and predicted damaging genetic variation leads to a reduction in MAPT expression, we asked if canonical loss-of-function in MAPT was depleted in AD cases vs. controls, but we did not detect a depletion of canonical loss-of-function variants, likely due to the small number of these qualifying variants (3 in cases, and 3 in controls).

## DISCUSSION

Many studies have been successful in reducing tau levels by targeting either tau modulators or knocking out tau itself.^11–13, 15, 86^ In mice, total tau ablation causes subtle motor deficits later in life; however, these deficits were not observed with partial tau reduction, which still provides therapeutic benefit.^11, 87^ Targeting tau through regulatory mechanisms controlling its expression may be a tractable method for reducing, instead of ablating, *MAPT* expression with limited off-target effects. Further, in modern genetics understanding how non-coding regions of the genome contribute to disease has been a major area of focus. Many methods have been developed to better identify candidate regulatory regions by assessing chromatin accessibility, modifications, and structure to determine interactions contributing to gene expression. However, each of these methods have limitations that lead to both false positives and negatives. By combining orthogonal genomics methods, we identified 94 *MAPT* candidate cis-regulatory elements and then sought to functionally interrogate these regions to determine which regulate *MAPT*. We tested all 94 regions using CRISPRi to determine their necessity for *MAPT* expression, and using 3’ RNA-seq, we determined if these regions were necessary for the expression of any other genes within the region. A subset of our nominated regions were selected for testing in a luciferase reporter assay to determine if these regions were sufficient for enhancer-like activity.

We identified two regions in the *FMNL1* locus as putative regulators of *MAPT* expression. We provide evidence in three independent chromatin conformation datasets that the region corresponding to 674,458 upstream of *MAPT* interacts with the *MAPT* promoter in excitatory and inhibitory neurons. While this region was not confirmed by RNA-seq, we observed significant reduction of *MAPT* expression by RT-qPCR. Importantly, we identified a novel regulator of *MAPT* in a region 20 kb downstream (Region −652,338) within exon 15 of *FMNL1*. A study by Birnbaum, et al.^88^ demonstrated that DNA sequences could act as a protein coding sequence in one tissue but regulate the expression of a nearby gene in another tissue. *FMNL1* is not expressed in any cell type in the DLPFC snRNA-seq data (data not shown) and is very lowly detected in the brain (gtexportal.org/home/gene/FMNL1). Therefore, this region could have dual function where it acts as a regulatory element of *MAPT* in tissues where the gene harboring the sequence is not expressed.

Moreover, this region was specifically linked to *MAPT* in nuclei from AD donors, and we found that this region is *∼* 1.5 kb downstream of a neuron-specific Capture-C interaction (chr17: 45,239,001-45,239,500) with the *MAPT* promoter. We confirmed using snATAC-seq from DLPFC tissue that this region was specifically accessible in excitatory and inhibitory neurons. This region was previously reported by Corces, et al. to have elevated interaction with the *MAPT* promoter by H3K27Ac HiChIP specifically in the H1 haplotype, which is associated with an increased risk for AD.^45^ Corces, et al. also provided single cell ATAC-seq evidence that this region harbors neuron-specific regulatory elements. While we only examined the function of this region in H1 haplotype neurons, we provide novel evidence that this region harbors an important CRE for *MAPT* regulation. Specific testing in H2 haplotype cell lines would be required to validate this cell type- and haplotype-specific regulation.

Within the *ARHGAP27* locus, we identified two regions as novel CREs of *MAPT*, corresponding to regions 464,677 and 461,949 upstream of *MAPT*. Both of these regions were identified as links to *MAPT* in both AD and control DLPFC tissues. The region 464,677 upstream of *MAPT* had enhancer-like activity in the luciferase reporter assay and showed significant knockdown of *MAPT* expression in CRISPRi experiments. Together these two lines of evidence prove that this region is a novel enhancer that regulates *MAPT* expression. Interestingly, both regions also showed a significant increase in *MAP3K14* expression upon targeting with CRISPRi. We also observe this inverse differential expression change when targeting a region within the first intron of *MAPT*-AS1 (Region −44,905). This region was also nominated by single nucleus multiomics and linked to several other genes, but not to *MAP3K14*. Additionally, in the RNA-seq analysis we observe that 19 regions tested significantly reduce MAP3K14 expression. *MAP3K14* is located over 500 kb upstream of *MAPT* near *FMNL1*. Several of these regions also show a trend of decreased *MAPT* expression that would suggest that they may be involved in *MAPT* regulation but not required for its expression. These observations suggest that there may be significant interaction between the *MAP3K14/FMNL1* region and *MAPT*, but this relationship requires further investigation.

Intronic CREs have been previously shown to regulate the expression of their host gene, and recently it was reported that intronic enhancers are enriched for tissue-specific activity.^89^ The proportion of intronic enhancer-like sequences was shown to be enriched in the most specialized cell types, like neurons, and they were shown to regulate genes involved with cell-type specific functions. We confirmed a CRE within the first intron of *MAPT* (Region 48,416). This region was linked to *MAPT* in both AD and control DLPFC tissues. Targeting this region with CRISPR dCas9-KRAB resulted in significant knockdown of both *MAPT* and *MAP3K14* expression, but it resulted in a significant increase in NSF expression. This intronic cCRE is particularly important because the H1c haplotype-tagging SNP rs242557 is located within this region. In our luciferase data, we tested the reference allele and did not observe a significant increase in luminescence signal. However, the alternate allele has been previously shown to increase luciferase activity when present and is associated with an increased risk of AD.^85^ Taken together, these results suggest that this specific CRE may have differing contributions to *MAPT* expression dependent on the haplotype context.

We observed a depletion of rare and predicted damaging genetic variation in AD cases in both nominated and confirmed CREs. While opposite to typical variant-burden studies, this effect direction would be expected under a model wherein damaging variation in these elements impairs enhancer function and thereby reduces expression of *MAPT*, which would likely be protective against AD. Thirty-one of the 41 identified genetic variants in CREs were annotated as non-coding. For the remaining 10 variants annotated as coding, all were annotated as coding for genes other than *MAPT*.

We speculate that the functional effect could still be regulatory for *MAPT*. For instance, we provide a high level of evidence for regulatory function of coding regions of *FMNL1* and *ARHGAP27* (**Figure 4**), which together account for 5 of the coding variants observed. The observation of depletion of rare and predicted damaging genetic variation in AD cases in both nominated and confirmed CREs is particularly important evidence that supports the value of detailed identification of CREs for AD-associated genes.

This study has several limitations. First, all experiments were performed in H1 haplotype cell lines. Therefore, regulatory regions identified here may not translate to the H2 haplotype, as the inversion of the locus may result in the loss of interaction of these regions with the *MAPT* promoter.^45^ Another limitation is that we performed CRISPRi experiments in a mixed culture of inhibitory and excitatory neurons and astrocytes. Given the variability in purity of this culture system, it is possible that there are false negatives for some neuron-specific regulatory regions. Future experiments examining these putative CREs in high-throughput CRISPR screens, like Perturb-seq,^90^ using pure neuronal cultures might reveal more neuron-specific regulatory regions. Additionally, due to the low-throughput nature of luciferase assays, we were not able to assess all regions for enhancer-like activity. Therefore, further investigation of these regions through methods like massively parallel reporter assays (MPRAs) executed in relevant cell types such as neurons are warranted to fully understand their function. While we focused on identifying CREs, specifically enhancers, there are several other non-coding regulatory mechanisms that are not assessed here. Furthermore, the focus on CRISPRi could lead to false negatives for regulatory regions that are not required for *MAPT* expression, but could still be involved if activated (which could be addressed with an alternative approach, CRISPRa), or that only have measurable effects on *MAPT* expression in the presence of specific alleles. A final limitation is that, due to the preference for an experimental design as a screen that is as comprehensive as possible, some experiments were limited in the number of experimental replicates.

## CONCLUSIONS

This study represents (to our knowledge) the most comprehensive evaluation to date of cis-regulatory elements important for expression of *MAPT*. Identification of these CREs not only facilitates better understanding how genetic variants around *MAPT* contribute to disease risk, but also provides important foundational knowledge of regulation of an important marker gene in the central nervous system. Future studies aimed at identifying TFs bound to these regulatory regions could point to new therapeutic targets, and because we have identified neuron-specific regulatory regions, drug screens targeting TFs bound to these regions could provide therapeutic targets with both target gene and cell-type specificity. This study also lays the groundwork for high-throughput approaches for fine mapping and/or combinatorial knockdown of CREs as well as for studies beyond the assessment conducted here on the effects of genetic variation in regulatory regions that are important for *MAPT* expression.

## LIST OF ABBREVIATIONS

AD: Alzheimer’s Disease

ADSP: Alxheimer’s Disease Sequencing Project

cCRE: candidate Cis-Regulatory Element

CRE: *Cis*-Regulatory Element

CBD: Corticobasal Degeneration

DEG: Differentially Expressed Gene

dELS: distal Enhance-Like Sequence

FDR: False Discovery Rate

FTD: Frontotemporal Dementia

GWAS: Genome Wide Association Study

LOAD: Late-Onset Alzheimer’s Disease

LD: Linkage Disequilibrium

MPRA: Massively Parallel Reporter Assay

Mb: Megabase

MAPT: Microtubule Associated Protein Tau

NPC: Neural Progenitor Cell

NFT: Neurofibrillary Tangle

OR: Odds Ratio

PD: Parkinson’s Disease

PCA: Principal Component Analysis

pELS: proximal Enhancer-Like Sequence

PLS: Promoter-Like Sequence

PSP: Progressive Supranuclear Palsy

qRT-PCR: Quantitative Reverse-Transcription

PCR SNP: Single Nucleotide Polymorphism

snRNA-seq: Single Nucleus RNA-sequencing

snATAC-seq: Single Nucleus ATAC-sequencing

TAD: Topologically Associated Domain

TF: Transcription Factor

TSS: Transcription Start Site

UMAP: Uniform Manifold Approximation and Projection

WNN: Weighted Nearest Neighbor

## DECLARIATIONS

### Availability of Data and Materials

The raw and processed data generated are available through NCBI GEO under series accession number GSE228121. All the code generated during this study is available at github.com/aanderson54/MAPT_cre. Variant data was obtained from the Alzheimer’s Disease Sequencing Project (ADSP) (NIAGADS accession number: NG00067).

## Acknowledgements

We thank Jacob M. Loupe for assisting A.G.A. with ChIP-seq analysis. This study was supported by BrightFocus fellowship A2019129F and NIH grants K99AG068271/5R00AG068271 awarded to J.N.C. as well as donors to the HudsonAlpha Foundation Memory and Mobility Program and Leo Fund.

## Author contributions

Conceptualization, J.N.C; Formal Analysis, A.G.A. and B.B.R.; Investigation, B.B.R., S.N.L., S.R., B.S.R., M.N.D., I.R., R.M.H., J.N.C., and E.B.; Data Curation, A.G.A. and B.B.R.; Writing–Original Draft, B.B.R., A.G.A., and J.N.C.; Writing– Review and Editing, L.F.R., J.N.C., S.J.C. and R.M.M.; Supervision, S.J.C., R.M.M., and J.N.C.; Funding, R.M.M. and J.N.C.

---

**Supplementary Fig. 1.**
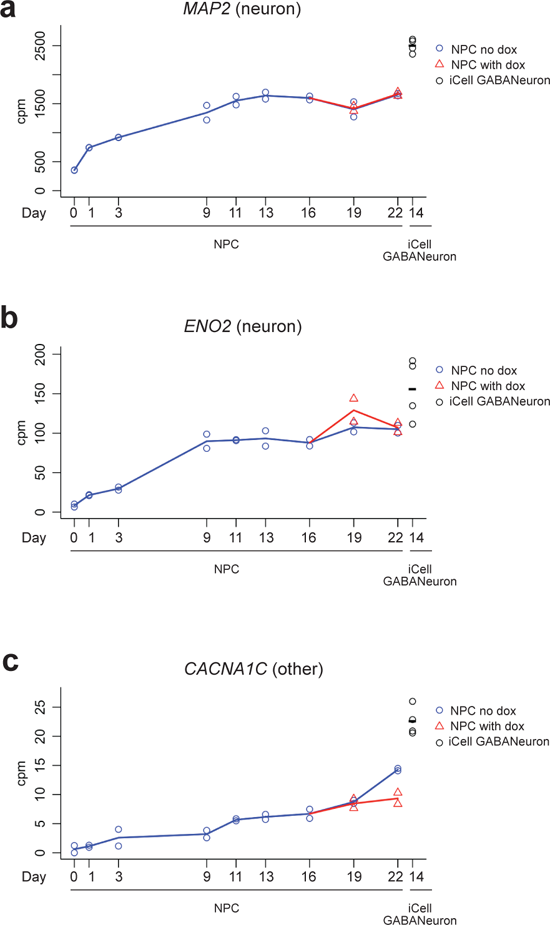
BrainPhys neuron differentiation time course for calculating differentation score, Related to Methods. Expression of key genes (**a.** *MAP2*, **b.** *ENO2*, and **c.** *CACNA1C*) used to determine level of differentiation of each replicate of differentiated neurons for CRISPRi experiments across 22 days compared to pure culture of inhibitory neurons (iCell GABANeuron day 14).

**Supplementary Fig. 2.**
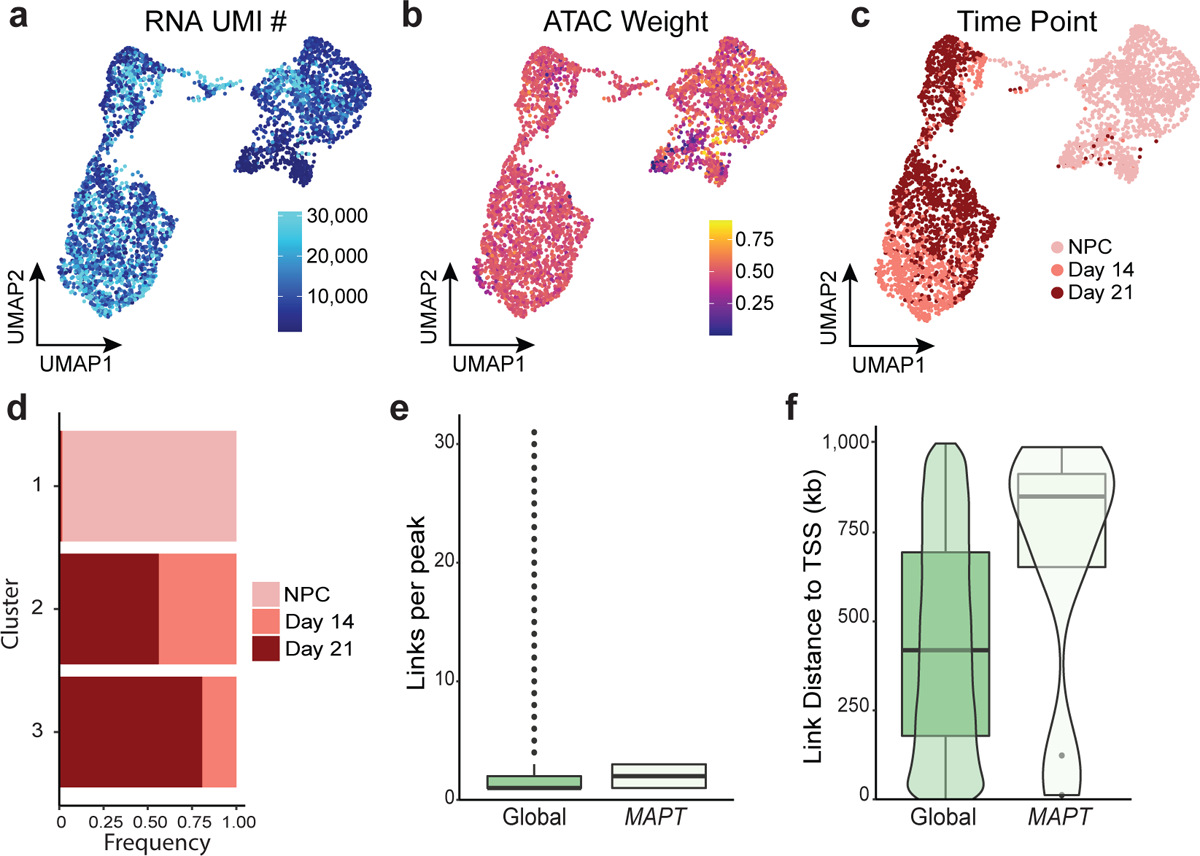
Integrating snRNA-seq and snATAC-seq and feature linkage description for iPSC-derived neuronal cultures, Related to Figure 1. **a.** WNN UMAP colored by RNA UMI counts. **b.** WNN UMAP colored by the proportion weight given to the snATAC-seq for creating the WNN graph. **c.** WNN UMAP colored by time point. **d.** The proportion of cells assigned to a cluster from each time point. **e.** Comparison of the distribution of links per ATAC peak of global link calls to *MAPT* links. **f.** Comparison of the distribution of the distance from linked ATAC peak to linked gene’s TSS for global links and *MAPT* links.

**Supplementary Fig. 3.**
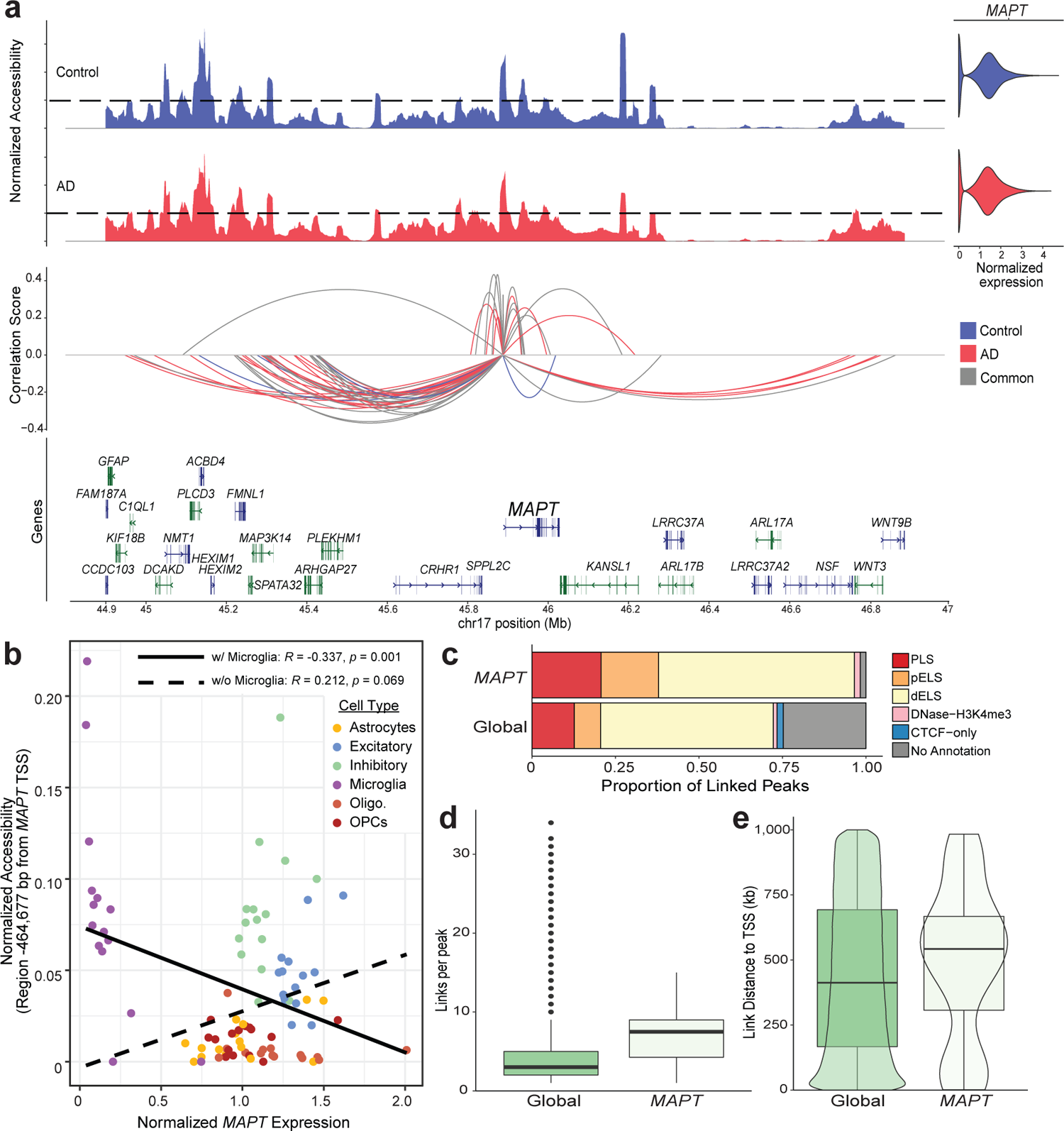
Nomination of cCREs using publicly available AD vs control single nucleus multiomics data. **a.** Linkage plot for all links to MAPT. Top panels: coverage plot of normalized accessibility (range 0 – 110,000) across all cell types separated by donor status (red = AD; blue = control). Bottom panel: significant AD and control peak-gene links. Arc height represents strength and direction of correlation. Arc color indicates if the link was AD-specific (red), control-specific (blue), or shared in both AD and control (”common”, gray). **b.** Example of negative correlation of accessibility (Target Region −464,677) with normalized MAPT expression by cell type. Solid line indicates correlation with microglia included. Dashed line indicates correlation with microglia cells removed. Spearman correlation and corresponding signficance are displayed with and without microglia. **c.** Comparison of proportion of linked peaks by ENCODE regualtory element annotation of global and *MAPT* links. **d.** Comparison of the distribution of links per ATAC peak of global links and *MAPT* links. **e.** Comparison of the distribution of the distance from linked ATAC peak to linked gene’s TSS of global links to *MAPT* links.

**Supplementary Fig. 4.**
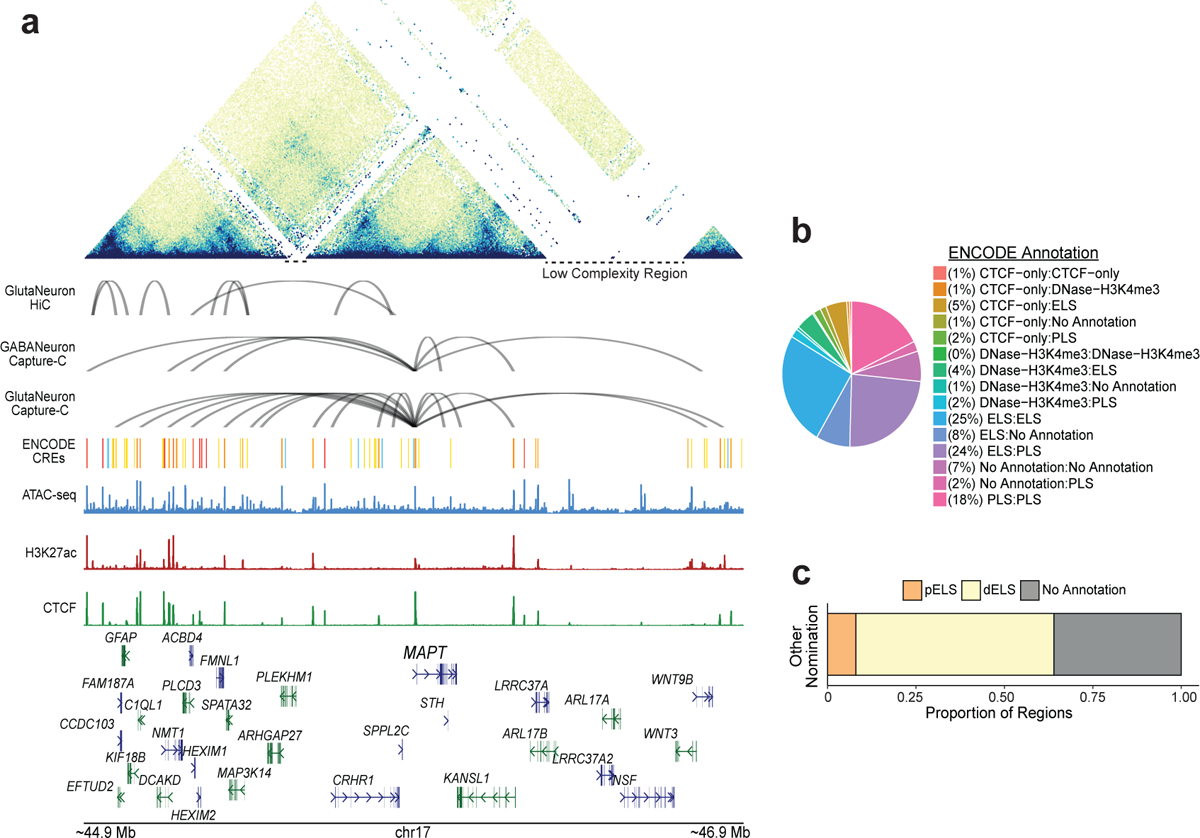
Description of chromatin annotation and structural analysis CREs, Related to Figure 2. **a.** Loop plot of HiCCUPs nominated loops in the *MAPT* locus. Top panel: Triangular plot showing TAD hierarchy. White spaces underlined with dashed lines represent regions of low complexity where sequences could not be adequately mapped. Middle panel: HiCCUPs loops in *MAPT* locus for HiC in iCell GlutaNeurons (top), Capture-C in iCell GlutaNeurons (middle), and Capture-C in iCell GABANeurons (bottom). Bottom panel: Track displaying ENCODE cCREs in the locus. ATAC-seq peaks and H3K27Ac and CTCF ChIP-seq peaks generated from iPSC-derived neuronal cultures differentiated for 14 days. **b.** Distribution of HiC loops by ENCODE annotation. **c.** Distribution of regions nominated from chromatin annotations and previously published data by ENCODE annotation.

**Supplementary Fig. 5.**
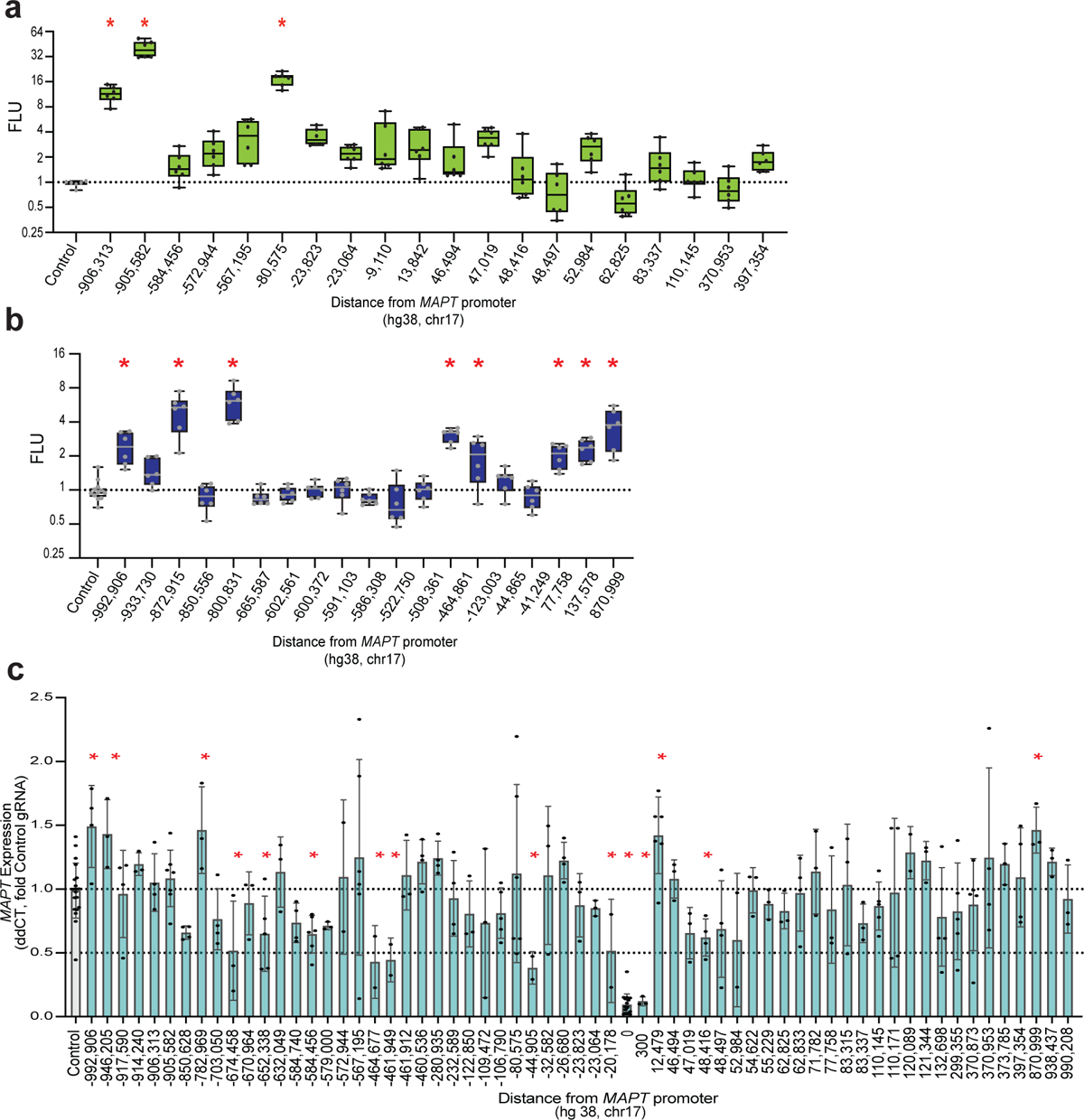
All functional validation, Related to Figure 4. **a.** Boxplots showing elements tested in luciferase assays in iCell GlutaNeurons (* p<0.05, ANOVA with Fisher’s LSD). **b.** Boxplots showing elements tested in KOLF2.1J-hNGN2 neurons (* p<0.05, ANOVA with Fisher’s LSD). **c.** RT-qPCR of *MAPT* expression. Barplots show regions tested in Day 14 BrainPhys differentiated neurons by CRISPRi experiments (*p <0.05, ANOVA with Fisher’s LSD).

## SUPPLEMENTARY INFORMATION

**Additional File 1:Supplementary Table 1.** Cluster DEGs from cultured neuron single nucleus multiomics before filtering low quality clusters. **Supplementary Table 2.** Significantly enriched GO terms from cultured neuron single nucleus multiomics before filtering low quality clusters. **Supplementary Table 3.** Cluster DEGs from cultured neuron single nucleus multiomics after filtering low quality clusters. **Supplementary Table 4.** Designed gRNAs.**Supplementary Table 5.** *MAPT* single nucleus multiomics links in cultured neurons. **Supplementary Table 6.** All linked genes for cultured neuron single nucleus multiomics *MAPT* linked peaks. **Supplementary Table 7.** *MAPT* single nucleus multiomics links in postmortem human brain. **Supplementary Table 8.** All linked genes for postmortem human brain single nucleus multiomics *MAPT* linked peaks. **Supplementary Table 9.** HiC *MAPT* HICCUPs called loops in iCell GlutaNeurons. **Supplementary Table 10.** iCell GlutaNeuron *MAPT* promoter Capture-C loops. **Supplementary Table 11.** iCell GABANeuron *MAPT* promoter Capture-C loops. **Supplementary Table 12.** “Other” nominated regions. **Supplementary Table 13.** Differential Capture-C loop analysis of NPCs and BrainPhys differentiated neurons. **Supplementary Table 14.** Differential Capture-C loop analysis of iCell GlutaNeurons and iCell GABANeurons. **Supplementary Table 15.** Regions selected for testing in luciferase assay. Supplementary Table 16. Regions overlapping publicly available functional genomics sets. **Supplementary Table 17.** RNA-seq differentially expressed genes. **Supplementary Table 18**. ADSP qualifying variants for burden analysis.

### Extended Acknowledgments

#### ADSP

The Alzheimer’s Disease Sequencing Project (ADSP) is comprised of two Alzheimer’s Disease (AD) genetics consortia and three National Human Genome Research Institute (NHGRI) funded Large Scale Sequencing and Analysis Centers (LSAC). The two AD genetics consortia are the Alzheimer’s Disease Genetics Consortium (ADGC) funded by NIA (U01 AG032984), and the Cohorts for Heart and Aging Research in Genomic Epidemiology (CHARGE) funded by NIA (R01 AG033193), the National Heart, Lung, and Blood Institute (NHLBI), other National Institute of Health (NIH) institutes and other foreign governmental and non-governmental organizations. The Discovery Phase analysis of sequence data is supported through UF1AG047133 (to Drs. Schellenberg, Farrer, Pericak-Vance, Mayeux, and Haines); U01AG049505 to Dr. Seshadri; U01AG049506 to Dr. Boerwinkle; U01AG049507 to Dr. Wijsman; and U01AG049508 to Dr. Goate and the Discovery Extension Phase analysis is supported through U01AG052411 to Dr. Goate, U01AG052410 to Dr. Pericak-Vance and U01 AG052409 to Drs. Seshadri and Fornage.

Sequencing for the Follow Up Study (FUS) is supported through U01AG057659 (to Drs. PericakVance, Mayeux, and Vardarajan) and U01AG062943 (to Drs. Pericak-Vance and Mayeux). Data generation and harmonization in the Follow-up Phase is supported by U54AG052427 (to Drs. Schellenberg and Wang). The FUS Phase analysis of sequence data is supported through U01AG058589 (to Drs. Destefano, Boerwinkle, De Jager, Fornage, Seshadri, and Wijsman), U01AG058654 (to Drs. Haines, Bush, Farrer, Martin, and Pericak-Vance), U01AG058635 (to Dr. Goate), RF1AG058066 (to Drs. Haines, Pericak-Vance, and Scott), RF1AG057519 (to Drs. Farrer and Jun), R01AG048927 (to Dr. Farrer), and RF1AG054074 (to Drs. Pericak-Vance and Beecham).

The ADGC cohorts include: Adult Changes in Thought (ACT) (U01 AG006781, U19 AG066567), the Alzheimer’s Disease Research Centers (ADRC) (P30 AG062429, P30 AG066468, P30 AG062421, P30 AG066509, P30 AG066514, P30 AG066530, P30 AG066507, P30 AG066444, P30 AG066518, P30 AG066512, P30 AG066462, P30 AG072979, P30 AG072972, P30 AG072976, P30 AG072975, P30 AG072978, P30 AG072977, P30 AG066519, P30 AG062677, P30 AG079280, P30 AG062422, P30 AG066511, P30 AG072946, P30 AG062715, P30 AG072973, P30 AG066506, P30 AG066508, P30 AG066515, P30 AG072947, P30 AG072931, P30 AG066546, P20 AG068024, P20 AG068053, P20 AG068077, P20 AG068082, P30 AG072958, P30 AG072959), the Chicago Health and Aging Project (CHAP) (R01 AG11101, RC4 AG039085, K23 AG030944), Indiana Memory and Aging Study (IMAS) (R01 AG019771), Indianapolis Ibadan (R01 AG009956, P30 AG010133), the Memory and Aging Project (MAP) (R01 AG17917), Mayo Clinic (MAYO) (R01 AG032990, U01 AG046139, R01 NS080820, RF1 AG051504, P50 AG016574), Mayo Parkinson’s Disease controls (NS039764, NS071674, 5RC2HG005605), University of Miami (R01 AG027944, R01 AG028786, R01 AG019085, IIRG09133827, A2011048), the Multi-Institutional Research in Alzheimer’s Genetic Epidemiology Study (MIRAGE) (R01 AG09029, R01 AG025259), the National Centralized Repository for Alzheimer’s Disease and Related Dementias (NCRAD) (U24 AG021886), the National Institute on Aging Late Onset Alzheimer’s Disease Family Study (NIA- LOAD) (U24 AG056270), the Religious Orders Study (ROS) (P30 AG10161, R01 AG15819), the Texas Alzheimer’s Research and Care Consortium (TARCC) (funded by the Darrell K Royal Texas Alzheimer’s Initiative), Vanderbilt University/Case Western Reserve University (VAN/CWRU) (R01 AG019757, R01 AG021547, R01 AG027944, R01 AG028786, P01 NS026630, and Alzheimer’s Association), the Washington Heights-Inwood Columbia Aging Project (WHICAP) (RF1 AG054023), the University of Washington Families (VA Research Merit Grant, NIA: P50AG005136, R01AG041797, NINDS: R01NS069719), the Columbia University Hispanic Estudio Familiar de Influencia Genetica de Alzheimer (EFIGA) (RF1 AG015473), the University of Toronto (UT) (funded by Wellcome Trust, Medical Research Council, Canadian Institutes of Health Research), and Genetic Differences (GD) (R01 AG007584). The CHARGE cohorts are supported in part by National Heart, Lung, and Blood Institute (NHLBI) infrastructure grant HL105756 (Psaty), RC2HL102419 (Boerwinkle) and the neurology working group is supported by the National Institute on Aging (NIA) R01 grant AG033193. The CHARGE cohorts participating in the ADSP include the following: Austrian Stroke Prevention Study (ASPS), ASPS-Family study, and the Prospective Dementia Registry-Austria (ASPS/PRODEM-Aus), the Atherosclerosis Risk in Communities (ARIC) Study, the Cardiovascular Health Study (CHS), the Erasmus Rucphen Family Study (ERF), the Framingham Heart Study (FHS), and the Rotterdam Study (RS). ASPS is funded by the Austrian Science Fond (FWF) grant number P20545-P05 and P13180 and the Medical University of Graz. The ASPS-Fam is funded by the Austrian Science Fund (FWF) project I904), the EU Joint Programme – Neurodegenerative Disease Research (JPND) in frame of the BRIDGET project (Austria, Ministry of Science) and the Medical University of Graz and the Steiermärkische Krankenanstalten Gesellschaft. PRODEM-Austria is supported by the Austrian Research Promotion agency (FFG) (Project No. 827462) and by the Austrian National Bank (Anniversary Fund, project 15435. ARIC research is carried out as a collaborative study supported by NHLBI contracts (HHSN268201100005C, HHSN268201100006C, HHSN268201100007C, HHSN268201100008C, HHSN268201100009C, HHSN268201100010C, HHSN268201100011C, and HHSN268201100012C). Neurocognitive data in ARIC is collected by U01 2U01HL096812, 2U01HL096814, 2U01HL096899, 2U01HL096902, 2U01HL096917 from the NIH (NHLBI, NINDS, NIA and NIDCD), and with previous brain MRI examinations funded by R01-HL70825 from the NHLBI. CHS research was supported by contracts HHSN268201200036C, HHSN268200800007C, N01HC55222, N01HC85079, N01HC85080, N01HC85081, N01HC85082, N01HC85083, N01HC85086, and grants U01HL080295 and U01HL130114 from the NHLBI with additional contribution from the National Institute of Neurological Disorders and Stroke (NINDS). Additional support was provided by R01AG023629, R01AG15928, and R01AG20098 from the NIA. FHS research is supported by NHLBI contracts N01-HC-25195 and HHSN268201500001I. This study was also supported by additional grants from the NIA (R01s AG054076, AG049607 and AG033040 and NINDS (R01 NS017950). The ERF study as a part of EUROSPAN (European Special Populations Research Network) was supported by European Commission FP6 STRP grant number 018947 (LSHG-CT-2006-01947) and also received funding from the European Community’s Seventh Framework Programme (FP7/2007-2013)/grant agreement HEALTH-F4- 2007-201413 by the European Commission under the programme “Quality of Life and Management of the Living Resources” of 5th Framework Programme (no. QLG2-CT-2002- 01254). High-throughput analysis of the ERF data was supported by a joint grant from the Netherlands Organization for Scientific Research and the Russian Foundation for Basic Research (NWO-RFBR 047.017.043). The Rotterdam Study is funded by Erasmus Medical Center and Erasmus University, Rotterdam, the Netherlands Organization for Health Research and Development (ZonMw), the Research Institute for Diseases in the Elderly (RIDE), the Ministry of Education, Culture and Science, the Ministry for Health, Welfare and Sports, the European Commission (DG XII), and the municipality of Rotterdam. Genetic data sets are also supported by the Netherlands Organization of Scientific Research NWO Investments (175.010.2005.011, 911-03-012), the Genetic Laboratory of the Department of Internal Medicine, Erasmus MC, the Research Institute for Diseases in the Elderly (014-93-015; RIDE2), and the Netherlands Genomics Initiative (NGI)/Netherlands Organization for Scientific Research (NWO) Netherlands Consortium for Healthy Aging (NCHA), project 050-060-810. All studies are grateful to their participants, faculty and staff. The content of these manuscripts is solely the responsibility of the authors and does not necessarily represent the official views of the National Institutes of Health or the U.S. Department of Health and Human Services.

The FUS cohorts include: the Alzheimer’s Disease Research Centers (ADRC) (P30 AG062429, P30 AG066468, P30 AG062421, P30 AG066509, P30 AG066514, P30 AG066530, P30 AG066507, P30 AG066444, P30 AG066518, P30 AG066512, P30 AG066462, P30 AG072979, P30 AG072972, P30 AG072976, P30 AG072975, P30 AG072978, P30 AG072977, P30 AG066519, P30 AG062677, P30 AG079280, P30 AG062422, P30 AG066511, P30 AG072946, P30 AG062715, P30 AG072973, P30 AG066506, P30 AG066508, P30 AG066515, P30 AG072947, P30 AG072931, P30 AG066546, P20 AG068024, P20 AG068053, P20 AG068077, P20 AG068082, P30 AG072958, P30 AG072959), Alzheimer’s Disease Neuroimaging Initiative (ADNI) (U19AG024904), Amish Protective Variant Study (RF1AG058066), Cache County Study (R01AG11380, R01AG031272, R01AG21136, RF1AG054052), Case Western Reserve University Brain Bank (CWRUBB) (P50AG008012), Case Western Reserve University Rapid Decline (CWRURD) (RF1AG058267, NU38CK000480), CubanAmerican Alzheimer’s Disease Initiative (CuAADI) (3U01AG052410), Estudio Familiar de Influencia Genetica en Alzheimer (EFIGA) (5R37AG015473, RF1AG015473, R56AG051876), Genetic and Environmental Risk Factors for Alzheimer Disease Among African Americans Study (GenerAAtions) (2R01AG09029, R01AG025259, 2R01AG048927), Gwangju Alzheimer and Related Dementias Study (GARD) (U01AG062602), Hillblom Aging Network (2014-A-004-NET, R01AG032289, R01AG048234), Hussman Institute for Human Genomics Brain Bank (HIHGBB) (R01AG027944, Alzheimer’s Association “Identification of Rare Variants in Alzheimer Disease”), Ibadan Study of Aging (IBADAN) (5R01AG009956), Longevity Genes Project (LGP) and LonGenity (R01AG042188, R01AG044829, R01AG046949, R01AG057909, R01AG061155, P30AG038072), Mexican Health and Aging Study (MHAS) (R01AG018016), Multi-Institutional Research in Alzheimer’s Genetic Epidemiology (MIRAGE) (2R01AG09029, R01AG025259, 2R01AG048927), Northern Manhattan Study (NOMAS) (R01NS29993), Peru Alzheimer’s Disease Initiative (PeADI) (RF1AG054074), Puerto Rican 1066 (PR1066) (Wellcome Trust (GR066133/GR080002), European Research Council (340755)), Puerto Rican Alzheimer Disease Initiative (PRADI) (RF1AG054074), Reasons for Geographic and Racial Differences in Stroke (REGARDS) (U01NS041588), Research in African American Alzheimer Disease Initiative (REAAADI) (U01AG052410), the Religious Orders Study (ROS) (P30 AG10161, P30 AG72975, R01 AG15819, R01 AG42210), the RUSH Memory and Aging Project (MAP) (R01 AG017917, R01 AG42210Stanford Extreme Phenotypes in AD (R01AG060747), University of Miami Brain Endowment Bank (MBB), University of Miami/Case Western/North Carolina A&T African American (UM/CASE/NCAT) (U01AG052410, R01AG028786), and Wisconsin Registry for Alzheimer’s Prevention (WRAP) (R01AG027161 and R01AG054047).

The four LSACs are: the Human Genome Sequencing Center at the Baylor College of Medicine (U54 HG003273), the Broad Institute Genome Center (U54HG003067), The American Genome Center at the Uniformed Services University of the Health Sciences (U01AG057659), and the Washington University Genome Institute (U54HG003079). Genotyping and sequencing for the ADSP FUS is also conducted at John P. Hussman Institute for Human Genomics (HIHG) Center for Genome Technology (CGT).

Biological samples and associated phenotypic data used in primary data analyses were stored at Study Investigators institutions, and at the National Centralized Repository for Alzheimer’s Disease and Related Dementias (NCRAD, U24AG021886) at Indiana University funded by NIA. Associated Phenotypic Data used in primary and secondary data analyses were provided by Study Investigators, the NIA funded Alzheimer’s Disease Centers (ADCs), and the National Alzheimer’s Coordinating Center (NACC, U24AG072122) and the National Institute on Aging Genetics of Alzheimer’s Disease Data Storage Site (NIAGADS, U24AG041689) at the University of Pennsylvania, funded by NIA. Harmonized phenotypes were provided by the ADSP Phenotype Harmonization Consortium (ADSP-PHC), funded by NIA (U24 AG074855, U01 AG068057 and R01 AG059716) and Ultrascale Machine Learning to Empower Discovery in Alzheimer’s Disease Biobanks (AI4AD, U01 AG068057). This research was supported in part by the Intramural Research Program of the National Institutes of health, National Library of Medicine. Contributors to the Genetic Analysis Data included Study Investigators on projects that were individually funded by NIA, and other NIH institutes, and by private U.S. organizations, or foreign governmental or nongovernmental organizations.

